# LRRC15 is an inhibitory receptor blocking SARS-CoV-2 spike-mediated entry *in trans*

**DOI:** 10.1101/2021.11.23.469714

**Authors:** Jaewon Song, Ryan D. Chow, Mario Pena-Hernandez, Li Zhang, Skylar A. Loeb, Eui-Young So, Olin D. Liang, Craig B. Wilen, Sanghyun Lee

## Abstract

SARS-CoV-2 infection is mediated by the entry receptor ACE2. Although attachment factors and co-receptors facilitating entry are extensively studied, cellular entry factors inhibiting viral entry are largely unknown. Using a *surface*ome CRISPR activation screen, we identified human LRRC15 as an inhibitory receptor for SARS-CoV-2 entry. LRRC15 directly binds to the receptor-binding domain (RBD) of spike protein with a moderate affinity and inhibits spike-mediated entry. Analysis of human lung single cell RNA sequencing dataset reveals that expression of LRRC15 is primarily detected in fibroblasts and particularly enriched in pathological fibroblasts in COVID-19 patients. *ACE2* and *LRRC15* are not co-expressed in the same cell types in the lung. Strikingly, expression of LRRC15 in ACE2-negative cells blocks spike-mediated viral entry in ACE2+ cell *in trans*, suggesting a protective role of LRRC15 in a physiological context. Therefore, LRRC15 represents an inhibitory receptor for SARS-CoV-2 regulating viral entry *in trans*.

## INTRODUCTION

Severe acute respiratory syndrome coronavirus 2 (SARS-CoV-2) is the causative agent of coronavirus disease 19 (COVID-19), representing a global health threat [1, 2]. SARS-CoV-2 belongs to the β-coronavirus family along with severe acute respiratory syndrome coronavirus (hereafter SARS-CoV-1) and middle east respiratory syndrome coronavirus (MERS-CoV) [3, 4]. Like SARS-CoV-1, SARS-CoV-2 utilizes angiotensin-converting enzyme 2 (ACE2) as a primary entry receptor [5, 6]. The viral structural protein spike (S), anchored on the surface of the viral envelope as homotrimers, binds to ACE2 and mediates the cellular entry of this virus [7]. The ectodomain of spike protein consists of the S1 and S2 subunits. The S1 subunit is comprised of the N-terminal domain (NTD) and the receptor binding domain (RBD) [8]. The RBD of spike protein directly binds to ACE2, which induces a conformational change that facilitates virus fusion either with endosomal membrane or with the plasma membrane [6, 9, 10]. This fusion event releases the SARS-CoV-2 genome into the cytoplasm [11, 12]. Importantly, the interaction between the RBD of spike and ACE2 is critical to determine several key features of SARS-CoV-2 infection. The high affinity interface between the RBD and ACE2 is associated with higher infectivity of SARS-CoV-2 compared to SARS-CoV-1 [13] and breaks a barrier of susceptibility of SARS-CoV-2 in murine hosts [14, 15]. Spike protein, specifically the RBD, is the primary target antigen for COVID vaccines in the market and interfering with the interface between RBD and ACE2 is the action mechanism of the majority of existing therapeutic antibodies [16], indicating the importance of RBD and its binding to the cellular receptor for controlling SARS-CoV-2.

Thus far, several cellular factors have been identified to facilitate cellular entry of SARS-CoV-2. However, it is unclear whether there are any cellular receptors that inhibit viral entry. Previous studies indicate that cleavage of spike protein by cellular proteases such as transmembrane protease serine 2 (TMPRSS2), cathepsins, and furin facilitates the entry of SARS-CoV-2 [9, 11, 17, 18]. Several cellular surface proteins or glycans facilitate viral entry by acting as an attachment factor, which includes neuropilin-1 [19, 20], heparan sulfate [21], and C-type lectins [22]. Alternative entry factors have been proposed such as AXL [23] and CD147 [24]. However, it remains to be elucidated whether any cellular entry factors regulate viral entry in a different manner.

In this study, we employed a screening method using the CRISPR activation (CRISPRa) technique. We generated a focused CRISPRa library, named *surface*ome, that covers ~6000 of all known/predicted surface proteins on the cellular plasma membrane. The *surface*ome screening with the SARS-CoV-2 spike protein revealed that human LRRC15 (leucin rich repeat containing 15) is a novel inhibitory receptor for SARS-CoV-2.

## RESULTS

### A *surface*ome CRISPR activation screen identified cellular receptors for spike protein of SARS-CoV-2

To identify host factors that regulate SARS-CoV-2 entry, we performed the *surface*ome CRISPRa screen and investigated which cellular proteins regulate spike binding to cells. We specifically selected approximately 6000 genes encoding plasma membrane proteins that contain either single or multiple transmembrane domains or are associated with the plasma membrane. We designed a CRISPR activation library consisting of four activating single guide RNAs (sgRNAs) per gene and 1,000 non-targeting control sgRNAs (**S1A Fig**). The screen was performed in a human melanoma cell line, A375, as this cell line does not express endogenous *ACE2* and does not interact with SARS-CoV-2 spike protein without ectopic expression of *ACE2* [21]. A375 cells containing catalytically “dead” Cas9 (dCas9) were transduced with the sgRNA library and selected to produce a pool of cells with induced expression of individual surface proteins. We measured the binding of Fc-tagged S1 subunit of SARS-CoV-2 spike to the cells by flow cytometry. Cells exhibiting high fluorescent signal intensity were sorted and subjected to genomic DNA extraction and sgRNA sequencing (**Fig 1A**). Two biologically independent screen results indicated two very distinct hits; ACE2 and LRRC15 (**Fig 1B**). ACE2 is reported to have a high affinity for the spike protein [6, 10]. LRRC15 is a leucin-rich repeat domain containing protein which is an orphan cancer-associated protein [25, 26]. There is no reported role of LRRC15 in SARS-CoV-2. An IgG isotype control and anti-CD45 staining identified IgG receptor genes (FCGR2C, FCGR3B) and CD45-encoding gene, PTPRC, as the top hit, respectively, confirming that the *surface*ome CRISPR screening efficiently identifies cellular receptors for targeted proteins (**Fig 1B, S1B Fig**).

**Fig 1.**
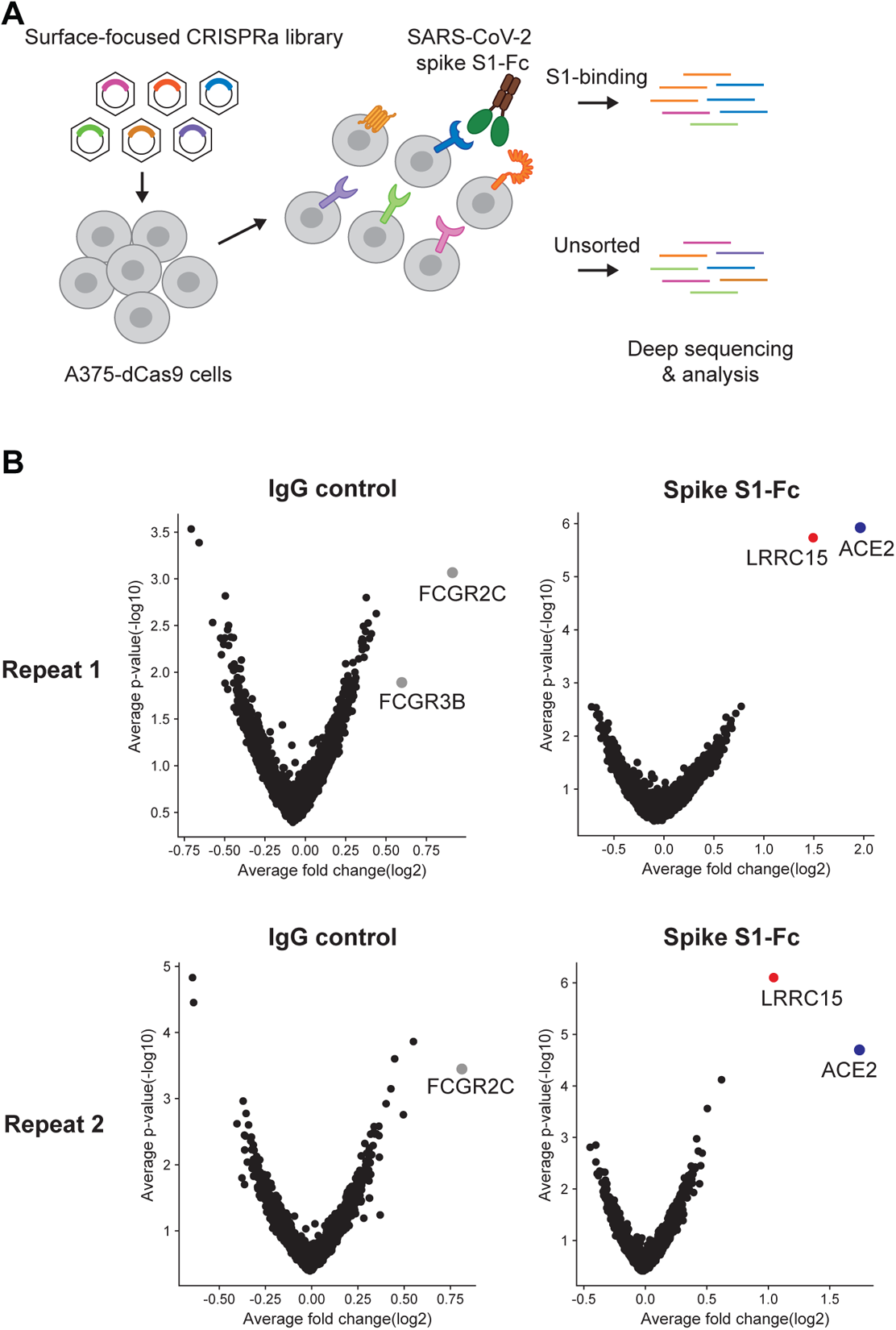
A *surface*ome-focused CRISPR activation screen identified cellular receptors binding with SARS-CoV-2 spike protein. (A) Schematic of a focused CRISPR activation screen for surface proteins interacting with SARS-CoV-2 spike S1-Fc fusion protein. (B) Volcano plots showing sgRNAs enriched or depleted in cells binding with SARS-CoV-2 spike S1-Fc or human IgG isotype control. Results from two biologically independent replicates are shown.

### LRRC15 directly interacts with the spike via the receptor-binding domain

To validate the screening result, we utilized two different human cell lines, A375 and HeLa. These two cell lines do not express endogenous *ACE2* and are not susceptible to SARS-CoV-2 without ectopic expression of *ACE2* [21, 27]. A375 and HeLa cells were transduced with two individual sgRNAs for *LRRC15* and a single sgRNA for *ACE2* to induce gene expression (**S2A-B Fig**). LRRC15-induced and ACE2-induced cells bound to the S1-Fc protein. The signal intensity in ACE2-induced cells was stronger than that of the LRRC15-induced cells. (**Fig 2A**). A similar pattern of protein-interaction was observed in HeLa cells (**Fig 2B, S2B Fig**).

**Fig 2.**
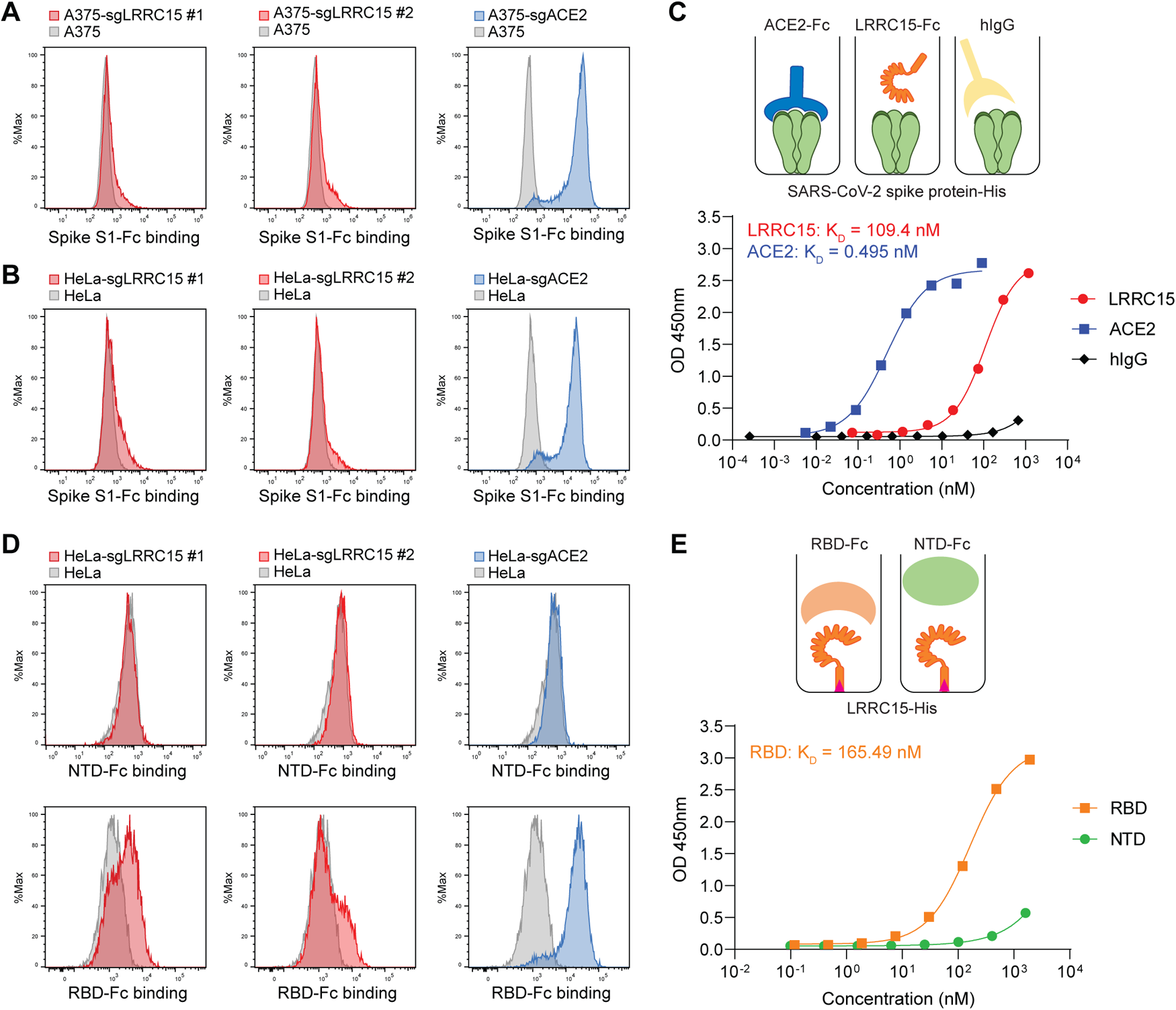
LRRC15 binds with SARS-CoV-2 spike protein at the receptor-binding domain. (A) A375 cells were transduced with indicated activating sgRNAs and incubated with SARS-CoV-2 spike S1-Fc fusion protein. Protein binding was measured by flow cytometry. (B) HeLa cells were transduced with indicated activating sgRNAs and incubated with SARS-CoV-2 spike S1-Fc fusion protein. Protein binding was measured by flow cytometry. (C) Dose-dependent binding of SARS-CoV-2 spike protein (Wuhan-Hu-1) to both ACE2 and LRRC15 with a Fc tag was determined by ELISA. Human IgG1 was included as a negative control. Dots indicate means of duplicates. (D) HeLa cells were transduced with indicated activating sgRNAs and incubated with SARS-CoV-2 spike NTD-Fc or RBD-Fc fusion protein. Protein binding was measured by flow cytometry. (E) The binding of the SARS-CoV-2 RBD and NTD to LRRC15 was measured by ELISA.

The interaction between LRRC15 and spike was further examined in a cell-free interaction model using recombinant proteins. An ELISA assay using recombinant LRRC15 and full-length spike indicated that LRRC15 directly interacts with the spike protein (K_D_ = 109 nM). The affinity between LRRC15 and the spike seems to be weaker than that of ACE2 and spike (**Fig 2C**). Interaction with spike proteins of different SARS-CoV-2 variants was confirmed. Recombinant full-length spike proteins of *α* (B.1.1.7), *β* (B.1.351), *ɣ* (P.1), *δ* (B.1.617.2), and *ι* (B.1.526) variants were tested and LRRC15 interacted with all of these spike proteins with similar affinity (**S2C Fig**).

ACE2 is known to interact with the spike protein via the RBD but does not interact with the NTD [28]. Interestingly, we identified that LRRC15 interacts with the spike in a similar way. Interaction assays in cells and in a cell-free assay using ELISA indicated that the RBD is sufficient to represent the interaction between LRRC15 and spike with a similar affinity compared to full-length S1 (**Fig 2D-E**). Next, we examined whether this interaction is specific to SARS-CoV-2 or conserved in other β coronaviruses. The ELISA assay using recombinant RBD protein of SARS-CoV-1 and MERS-CoV showed that LRRC15 binds to spike of SARS-CoV-1 with similar affinity but does not interact with spike of MERS-CoV (**S2D Fig**). These results indicate that LRRC15 is a novel cellular binding protein for the spike protein of SARS-CoV-1 and -2 and directly interacts with the spike via the RBD.

### LRRC15 suppresses entry of SARS-CoV-2

We next investigated whether LRRC15 regulates the entry process of SARS-CoV-2. Pseudotyping a heterologous viral envelope with spike protein has been utilized to study the entry process of SARS-CoV-2 [29, 30]. To monitor viral entry, we utilized a replication-incompetent VSV pseudovirus system that harbors the spike protein of SARS-CoV-2 on the viral envelope and expresses GFP reporter [31]. The LRRC15-induced A375 or HeLa cells were infected with the pseudovirus and the infectivity was monitored by flow cytometry. Wildtype control A375 and HeLa cells were not susceptible to the spike-pseudotyped virus, consistent with previous reports [21, 27] (**S3A-B Fig**). ACE2 but not LRRC15 expression was sufficient to support viral entry. The VSV-G-coated pseudovirus entered into all tested cell lines with a similar efficiency (**S3A-B Fig**). These data suggest that LRRC15 does not function as an entry receptor for SARS-CoV-2.

To examine whether LRRC15 regulates ACE2-mediated viral entry, we first generated a clonal HeLa cell line stably expressing ACE2, designated as HeLa-ACE2, and confirmed the high surface expression of ACE2 and susceptibility to SARS-CoV-2 spike-pseudovirus (**S3C-D Fig**). Using this cell line, we induced LRRC15 expression with two different sgRNAs and measured the efficiency of pseudoviral entry. The increased LRRC15 expression resulted in a significant decrease in the spike-pseudotyped VSV entry, while the induction of an unrelated gene, CD45, showed similar infectivity compared to the mock control (**Fig 3A, S3E Fig**). The entry of VSV-G-pseudotyped virus was not affected by LRRC15 gene induction, indicating the LRRC15-mediated inhibition is specific to the entry of SARS-CoV-2. The inhibitory function of LRRC15 was confirmed by ectopic overexpression of its cDNA as well (**Fig 3B, S3F Fig**). Of note, a larger reduction in viral entry was observed with the sgRNA-mediated gene induction, likely due to its higher gene expression of LRRC15 on cells compared to the cDNA overexpression (**S3E-F Fig**). The suppressed viral entry was observed in other pseudotyped viruses containing the spike of multiple variants of SARS-CoV-2 (i.e., α, β, ɣ, and δ) and SARS-CoV-1 (**Fig 3C-D**), as expected by their similar binding affinities with LRRC15 (**S2C-D Fig**). Therefore, these data indicate that LRRC15 suppresses spike-mediated viral entry, and the binding and suppression activity are specific to SARS-CoV-2 and SARS-CoV-1.

**Fig 3.**
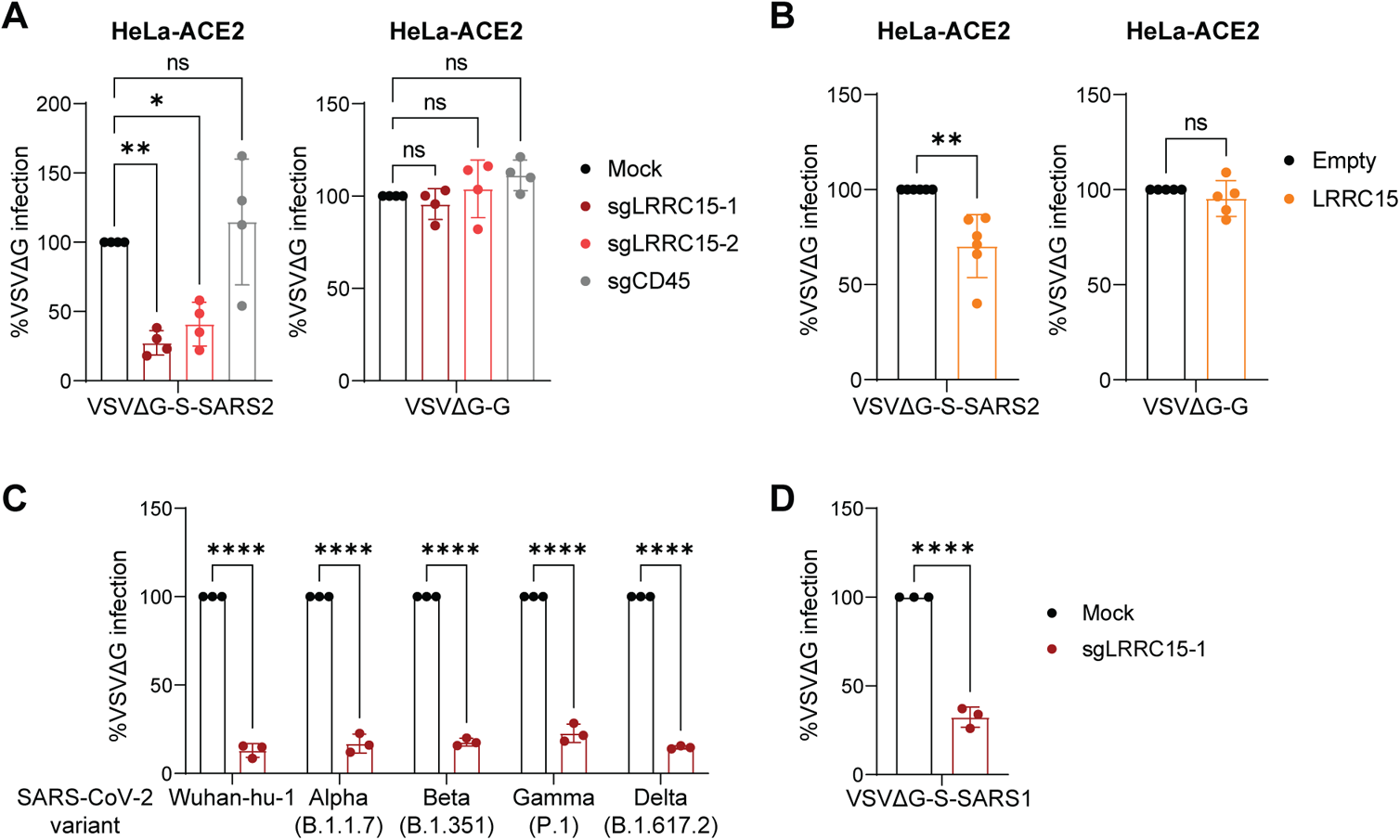
LRRC15 inhibits ACE2-mediated SARS-CoV-2 entry. (A) HeLa-ACE2 cells were transduced with indicated activating sgRNAs and infected with VSV pseudoviruses, VSVΔG-S-SARS2 or VSVΔG-G. GFP signal was measured at 20 hpi by flow cytometry and normalized to mock control (n=4). (B) HeLa-ACE2 cells expressing LRRC15 or empty vector were infected with VSV psuedoviruses and GFP signal was measured at 20 hpi by flow cytometry normalized to empty control (n=6 for VSVΔG-S-SARS2 and n=5 for VSVΔG-G). (C) LRRC15-induced or mock HeLa-ACE2 cells were infected with VSV pseudoviruses harboring spike proteins of different SARS-CoV-2 variants. GFP signal was measured at 20 hpi by flow cytometry (n=3). (D) LRRC15-induced or mock HeLa-ACE2 cells were infected with VSV pseudoviruses harboring SARS-CoV-1 spike. GFP signal was measured at 20 hpi by flow cytometry (n=3). Data represent means ± SEM (A-D). Data were analyzed by one-way ANOVA with Dunnett’s multiple comparisons test (A) or unpaired two-tailed t test (B-D). ns, not significant; *p < 0.05; **p < 0.01; ****p < 0.0001.

### LRRC15 accumulates cell-attached viruses on the membrane and does not compete with ACE2

As SARS-CoV-2 entry is primarily dependent on ACE2, we assessed whether LRRC15 alters protein expression of ACE2. In HeLa-ACE2 cells, the level of ACE2 surface expression was unaltered by either sgRNA-mediated gene induction or ectopic cDNA expression of LRRC15 (**Fig 4A**). Interestingly, we found that spike-coated viruses were sequestered on the cellular surface of LRRC15-expressing cells. In the attachment assay, we measured the attachment to cells by incubating spike-pseudotyped viruses and cells on ice, allowing attachment on the cell membrane and preventing internalization of viruses. As expected, the binding of SARS-CoV-2 spike-pseudotyped viruses to cells was dependent on ACE2 expression and the binding was not altered by inducing a control gene, CD45. Importantly, viruses were highly accumulated in LRRC15-induced HeLa-ACE2 cells (i.e., 3 to 5.5-fold increases in viral copies) compared to the control (**Fig 4B**). Preincubation of soluble LRRC15 protein with spike-pseudotyped viruses partly blocked the viral entry in HeLa-ACE2 cells at high concentration, while preincubation with soluble ACE2 completely blocked the viral entry (**Fig 4C**), suggesting that the inhibitory function of LRRC15 may require localization to the cellular membrane and/or the cytosolic domain of LRRC15 for its functional efficiency. In summary, these results suggest that LRRC15 inhibits SARS-CoV-2 entry by restricting the internalization of virions into the cell through binding to the spike protein on the viral envelope.

**Fig 4.**
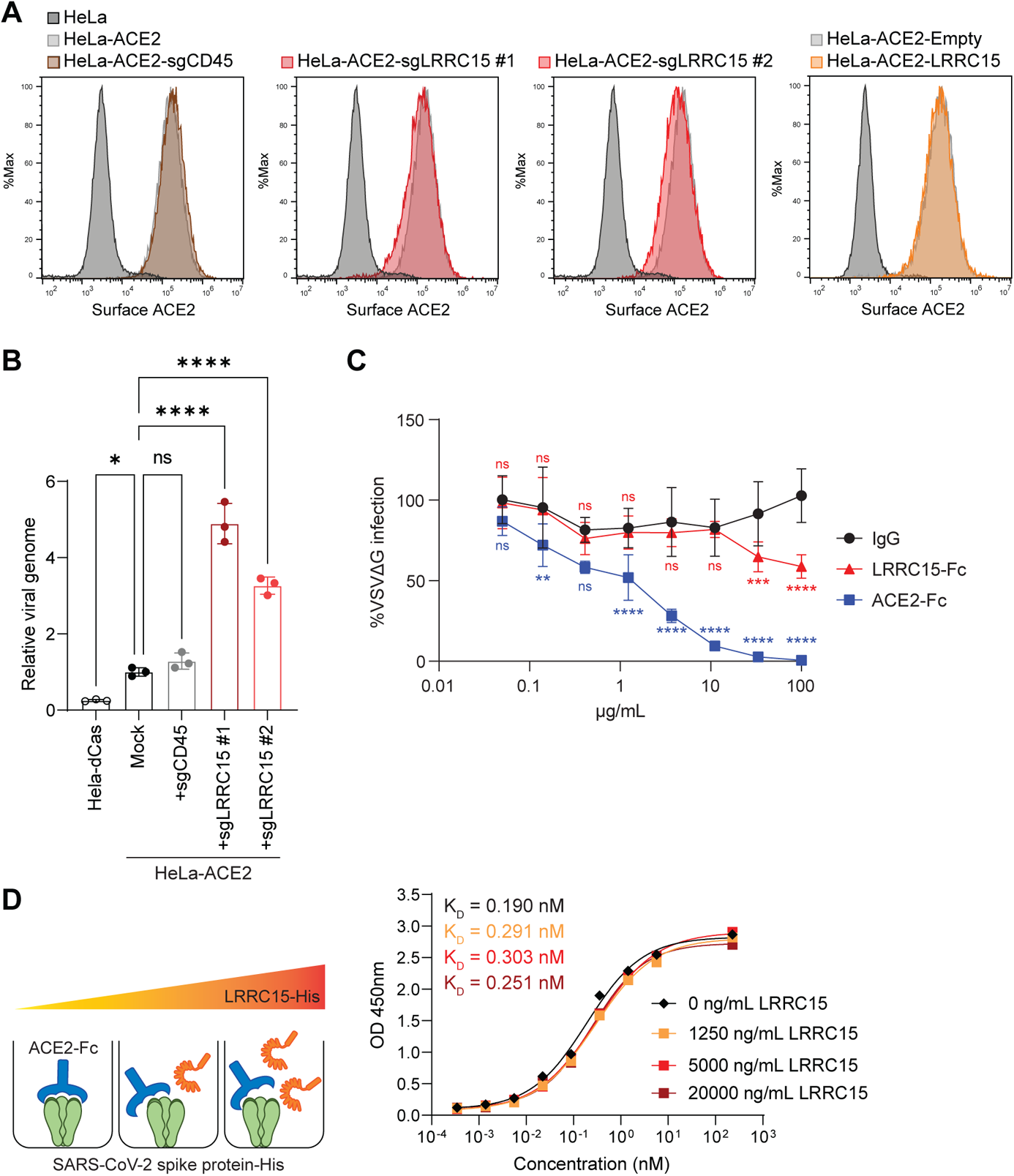
LRRC15 enhances SARS-CoV-2 attachment to the surface of ACE2-expressing cells. (A) Cell surface expression of ACE2 was measured in HeLa-ACE2 cells transduced with indicated activating sgRNAs or a LRRC15-expressing vector. (B) HeLa or HeLa-ACE2 cells transduced with indicated activating sgRNAs were incubated with VSVΔG-S-SARS2 for 1 h on ice and washed three times with cold cell culture media. Viral genome copies were quantified by RT-qPCR and normalized to HeLa-ACE2 cells (n=3). (C) VSVΔG-S-SARS2 were incubated with ACE2-Fc, LRRC15-Fc, or IgG control for 1 h, prior to inoculating HeLa-ACE2 cells. Viral infectivity was quantified by measuring GFP signal at 20 hpi by flow cytometry and normalized to no antibody control (n=6). Statistical significance was determined compared to IgG control at each dilution. (D) Competition assays between Fc-tagged ACE2 and His-tagged LRRC15 for immobilized His-tagged SARS-CoV2 spike protein. Premixture of LRRC15-His at 4 different concentrations with a dilution series of ACE2-Fc was added and anti-human HRP determined the amount of ACE2-Fc remaining in the presence of competitor LRRC15-His through a colorimetric readout. Data represent means ± SEM (B, C). Data were analyzed by one-way ANOVA (B) or two-way ANOVA (C) with Dunnett’s multiple comparisons test. ns, not significant; *p < 0.05; **p < 0.01; ***p < 0.001; ****p < 0.0001.

To test whether LRRC15 directly binds to ACE2, we utilized His-tagged-LRRC15, SARS-CoV-2 spike protein, and MERS-CoV spike protein and assessed their interactions with Fc-tagged ACE2. Recombinant ACE2 did not show detectable binding to recombinant LRRC15 protein or the MERS-CoV spike whereas binding to SARS-CoV-2 spike was confirmed with high affinity (**S4A Fig**) [32]. Since both ACE2 and LRRC15 bind to the RBD of SARS-CoV-2 spike, we investigated whether LRRC15 competes with ACE2 for binding on the spike protein. The interaction between ACE2 and SARS-CoV-2 spike was measured in the presence of recombinant LRRC15 protein. Even at high concentrations, LRRC15 did not affect the spike-ACE2 binding (**Fig 4D**). We confirmed SARS-CoV-2 spike S1-Fc binding to Hela-ACE2 cells was not altered by gene-induction or ectopic expression of LRRC15 (**S4B Fig**). Taken together, ACE2-mediated viral entry of SARS-CoV-2 is suppressed by LRRC15 on the cell membrane through its direct binding to the RBD without competition between LRRC15 and ACE2.

### LRRC15 is expressed in distinct cell types in human lung and is associated with pathological fibroblasts

To better understand the function of LRRC15 in a physiologic context, we first explored its expression in human lung samples unaffected by SARS-CoV-2. Two independent single-cell RNA-sequencing (scRNA-seq) datasets of non-COVID-19 human lungs [33, 34] revealed that *LRRC15* was predominantly expressed in a subset of fibroblasts and lymphatic endothelial cells (**S5A-B Fig**). For instance, in the Tissue Stability Cell Atlas dataset, a significant proportion of fibroblasts and lymphatic endothelial cells expressed *LRRC15* (**S5B Fig**). Of note, the cell types that expressed *LRRC15* did not co-express *ACE2*.

Having defined fibroblasts and lymphatic endothelial cells as the main cell types in the lung that express *LRRC15*, we sought to explore if any clinical features were associated with *LRRC15* expression in the lung. Utilizing the large cohort of human lung RNA-seq samples from the Genotype-Tissue Expression project (GTEx) [35], we constructed a multivariable regression model between *LRRC15* expression and various clinical factors as predictors. Specifically, we included age, sex, diabetes (type 1 or type 2), hypertension, body mass index (BMI), smoking, and ventilator status at time of death as independent variables in the model. We observed that *LRRC15* expression was not significantly associated with age, sex, hypertension, BMI, smoking, or type 1 diabetes (**S5C Fig**). Strikingly, *LRRC15* expression was significantly decreased in patients that were on a ventilator prior to death (**S5D Fig**). We note that the causality of this association cannot be ascertained due to the retrospective nature of the dataset: it is unclear whether ventilator usage leads to lower *LRRC15* expression, or whether patients with conditions that subsequently require mechanical ventilation have baseline alterations in lung physiology associated with decreased *LRRC15* expression.

In order to gain further insight on *LRRC15* expression in the lung, we calculated the correlation between *LRRC15* and all other genes in the GTEx lung dataset. We observed that genes such as *SOX4*, *FRMD6*, *FAP*, *ENAH*, *PRRX1*, *CD200*, and *VCAM1* were positively correlated with *LRRC15* (**S5E Fig**). In contrast, *ACE2* was negatively correlated with *LRRC15* (**S5F Fig**), which is consistent with their distinct cell type-specific expression patterns (**S5A-B Fig**). We subsequently mapped these highly correlated genes to specific lung cell types, finding that fibroblasts and lymphatic endothelial cells also expressed several of the genes that were positively correlated with *LRRC15* (**S5G Fig**). We further observed that *MAOA*, which showed the strongest negative correlation with *LRRC15* in the bulk lung RNA-seq cohort, was mostly expressed in alveolar type 2 cells (**S5E Fig**).

To explore the clinical relevance of LRRC15 to COVID-19 pathophysiology, we analyzed two independent scRNA-seq datasets of lungs from deceased COVID-19 patients [36, 37]. Consistent with our analyses of non-COVID-19 lungs, we found that fibroblasts and lymphatic endothelial cells had the highest levels of *LRRC15* expression (**Fig 5A-B**). Interestingly, we observed that *LRRC15* expression was particularly enriched in the pathological fibroblast subpopulation. This recently identified fibroblast subset (defined by high *CTHRC1* expression) has been implicated as a key contributor to idiopathic pulmonary fibrosis [38] and may also drive lung fibrosis in COVID-19 patients [36]. Indeed, the relative proportion of pathological fibroblasts and intermediate pathological fibroblasts was significantly increased in COVID-19 patients compared to controls (**Fig 5C**). Further supporting the association between *LRRC15* and disease-associated fibroblast cell states, there was a progressive gradient of *LRRC15* expression from alveolar fibroblasts (0.26%) to intermediate pathological fibroblasts (2.61%), and finally to pathological fibroblasts (4.52%).

**Fig 5.**
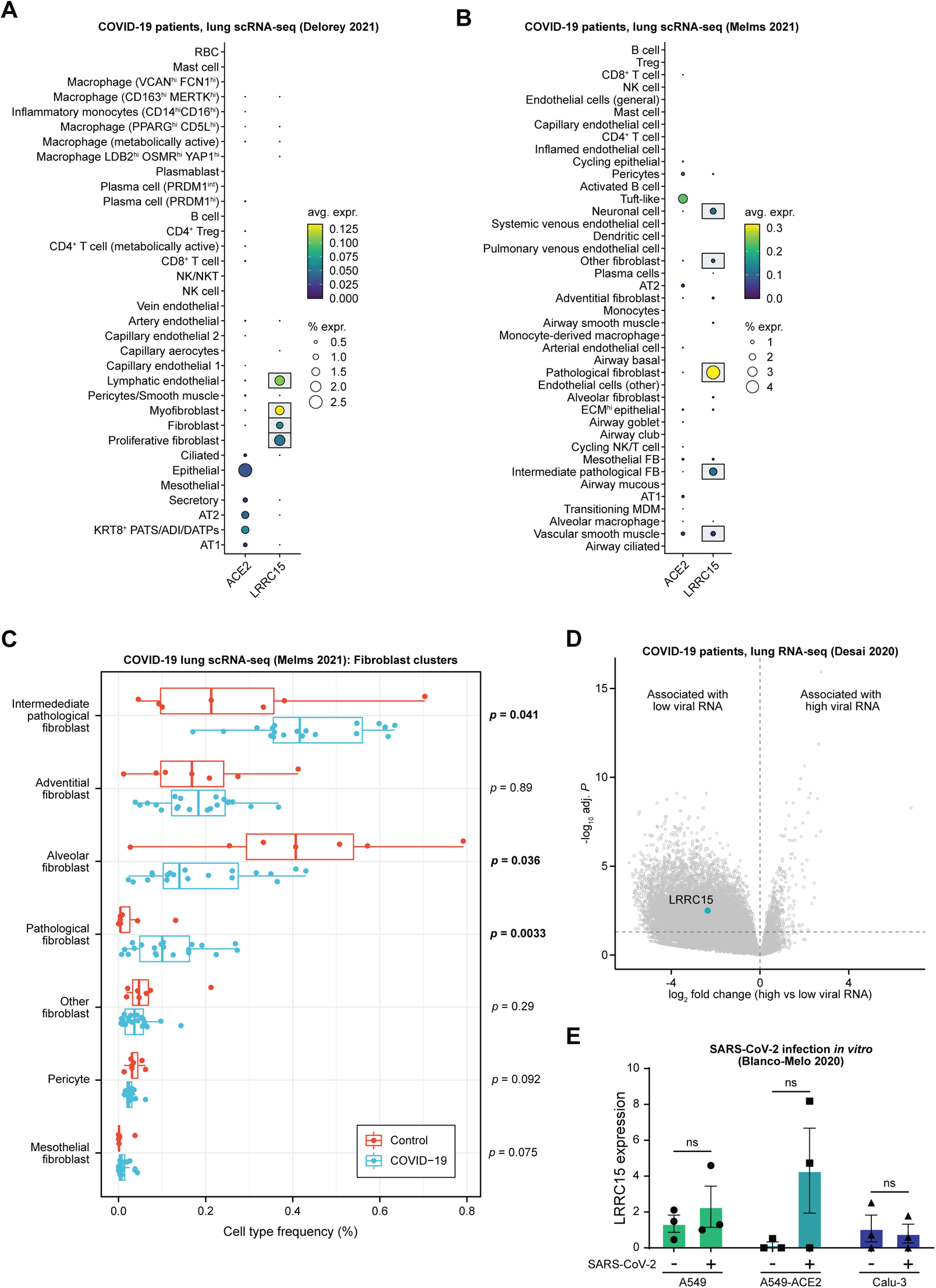
LRRC15 expression is enriched in the fibroblasts of COVID-19 patients and associated with reduced SARS-CoV-2 viral burden. (A-B) Cell type-specific expression of *ACE2* and *LRRC15*, assessed by scRNA-seq of lungs from deceased COVID-19 patients (A) Delorey et al, 2021 [37] or (B) Melms et al, 2021 [36]. (C) Boxplots of the relative frequencies of fibroblast subtypes among total fibroblasts, comparing COVID-19 patients (blue) to non-COVID-19 controls (red). Statistical significance was assessed by two-tailed unpaired Mann-Whitney test. (D) Volcano plot of differentially expressed genes in the lungs of deceased COVID-19 patients, comparing samples with high vs low SARS-CoV-2 RNA levels at the time of death [39]. Genes with positive log_2_ fold changes are associated with high viral burden, while genes with negative log_2_ fold changes are associated with low viral burden. (E) *LRRC15* expression in lung cell lines (A549, A549-ACE2, Calu-3), comparing mock controls vs SARS-CoV-2 infected samples. Statistical significance was assessed by two-tailed unpaired Welch’s t-test.

Given that our experiments pointed to LRRC15 as an antiviral restriction factor, we sought to specifically investigate whether *LRRC15* is associated with viral burden and disease progression in COVID-19 patients. We analyzed a bulk RNA-seq dataset of lungs from COVID-19 patients [39] with high or low SARS-CoV-2 viral burden at the time of autopsy. While patients with high vs low viral burden might correspond to two distinct disease phenotypes, patients with high viral RNA load had experienced shorter duration of illness before death, suggesting that these patients had failed to control the virus and died during the acute phase of infection [39]. On the other hand, patients with low viral RNA load had longer duration of illness before death, consistent with a scenario in which they had successfully controlled the virus but subsequently died from sequalae of the infection. With this framework in mind, we found that expression of *LRRC15* was significantly higher in patients with low SARS-CoV-2 viral burden (**Fig 5D**). Though the causality underlying this relationship is unclear, it is supportive of the antiviral function of LRRC15 that we have identified here. Of note, SARS-CoV-2 infection of lung epithelial cell lines *in vitro* did not lead to significant changes in *LRRC15* expression (**Fig 5E**), which is consistent with our prior analyses pointing to a fibroblast-specific role of *LRRC15* in COVID-19. Collectively, our analyses point to a model in which SARS-CoV-2 infection induces the emergence of a pathological fibroblast state with increased LRRC15 expression.

### LRRC15 in ACE2-negative cells inhibits SARS-CoV-2 infection *in trans*

Since the scRNA-seq analysis revealed that *LRRC15* is not co-expressed with *ACE2* within the same cell types in the lung, we hypothesized that LRRC15 inhibits SARS-CoV-2 entry into ACE2-positive cells *in trans*. To test this hypothesis, HeLa-ACE2 cells were co-cultured with either HeLa-control or HeLa-sgLRRC15 cells (i.e., ACE2-negative cells) and were subsequently infected with spike-pseudotyped viruses (**Fig 6A**). The GFP reporter signal was only detected in the ACE2+ cells, confirming LRRC15 alone does not permit viral entry (**S6A Fig**). Compared to the control HeLa cells, co-culturing with HeLa-sgLRRC15 cells resulted in a significant reduction of viral entry in ACE2+ cells when co-cultured at 1:4 ratio (**Fig 6B**). At two different titers of viral infection, the similar pattern of reduction was observed. The same trend of *trans*-inhibition was confirmed by the spike-pseudotyped viral infection with the delta (B.1.617.2) variant (**Fig 6C**). Co-culture of HeLa-ACE2 cells and HeLa-sgLRRC15 cells at 1:1 ratio exhibited significant restriction activities, although the magnitude of suppression is slightly weaker as expected (**S6B-C Fig**). Immunofluorescence staining with the co-culture model further confirmed the spike binding on LRRC15+ cells and strong co-localization of LRRC15 and spike on these cells (**Fig 6D**). Spike-pseudotyped virus-infected cells exhibited speckle-like staining patterns for spike. The co-culture condition with HeLa-sgLRRC15 showed that a significant proportion of spike speckles were detected on LRRC15+ cells. Most interestingly, spike speckles on the co-culture condition with HeLa-sgLRRC15 cells retained overtime without further entry progress, whereas the spike speckles rapidly disappeared in cells within 60 min in the co-culture condition with wildtype HeLa cells (**Fig 6E**). These results indicate that spike-binding on LRRC15+ cells is not functional binding for entry, but for non-infectious sequestration of virions. Collectively, these results highlight the *trans*-inhibitory function of LRRC15 proven using a co-culture model, suggesting that expression of LRRC15 in SARS-CoV-2-non-permissive fibroblasts protects permissive cells against viral infection in the lung (**Fig 6F**).

**Fig 6.**
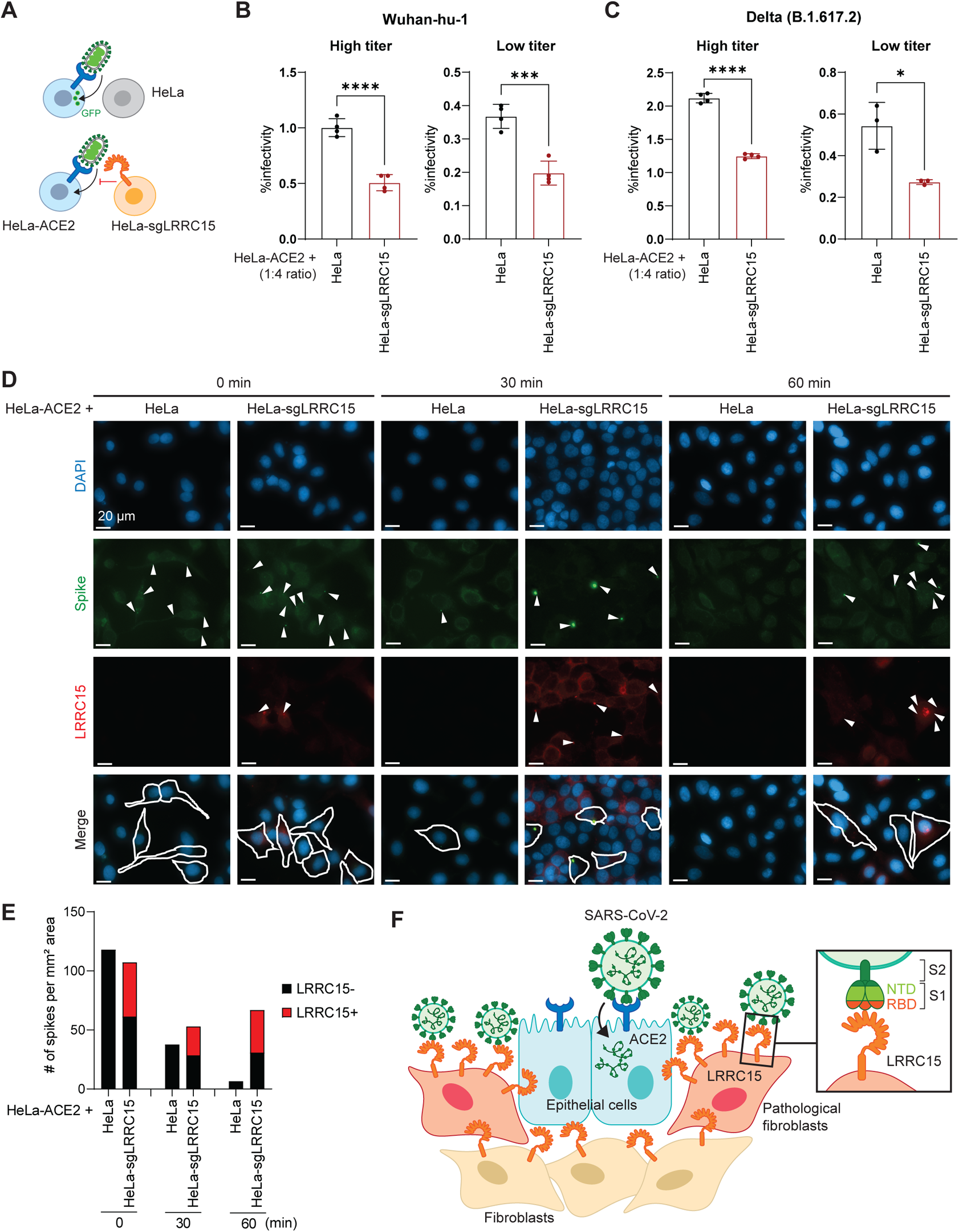
LRRC15 inhibits ACE2-mediated SARS-CoV-2 entry *in trans*. (A) Schematic of trans-inhibition assay with ACE2+ and LRRC15+ cells. (B) HeLa-ACE2 cells were co-cultured with HeLa or HeLa-sgLRRC15 cells at 1:4 ratio and infected with VSV pseudovirus harboring spike of SARS-CoV-2 Wuhan-hu-1 strain at high or low titer. GFP signal was measured at 20 hpi by flow cytometry (n=4). Representative of three independent experiments are shown. (C) Trans-inhibition assay was performed as in (B) with VSV pseudovirus harboring spike of SARS-CoV-2 Delta variant (B.1.617.2 strain) (n=4). Representative of three independent experiments are shown. (D) Representative images of immunofluorescence staining of SARS-CoV-2 spike (green), LRRC15 (red), and DAPI (blue). HeLa-ACE2 cells were co-cultured with HeLa or HeLa-sgLRRC15 cells, inoculated with VSVΔG-S-SARS2 for 1 h on ice and incubated at 37°C to allow internalization. Cells were harvested at indicated timepoints and subjected to staining. The white arrowheads indicate spikes. The scale bar indicates 20 µm. (E) Quantification was performed by calculating the number of spikes on LRRC15- or LRRC15+ cells per mm^2^ area from multiple images per sample. (F) Proposed working model of the LRRC15-mediated inhibition of SARS-CoV-2 entry. Data represent means ± SEM and were analyzed by unpaired two-tailed t test (B, C). *p < 0.05; ***p < 0.001; ****p < 0.0001.

## DISCUSSION

In this study, we employed a *surface*ome CRISPRa library to identify cellular receptors for SARS-CoV-2 by staining cells with a recombinant spike protein. The *surface*ome screening revealed two distinct hits: ACE2, the *bona fide* entry receptor, and LRRC15, a novel inhibitory receptor. Binding assays using recombinant proteins in cells and in cell-free models showed that LRRC15 directly interacts with the RBD of spike with a moderate affinity (*K_D_* = 43~148 nM, depending on domain and variant). Although ACE2 also interacts with the spike via the RBD, the interaction of LRRC15-RBD does not compete with the interaction of ACE2-RBD or stabilize it either. Further studies will be necessary to determine the interface of LRRC15-RBD at a higher resolution. Most strikingly, LRRC15 inhibited the spike-mediated viral entry not only in the same cells, but also in neighboring cells *in trans*, providing a unique concept of viral entry inhibition by an inhibitory receptor.

The inhibition of viral entry by LRRC15 represents a direct effector by interacting with the spike protein itself. The leucin-rich repeat (LRR) domain is functionally linked with sensing of pathogen-associated molecular patterns (PAMPs) in a number of cases [40]. The LRR domain proteins are highly conserved throughout evolution including in plants, providing a prominent protection against pathogens [41, 42]. Given the role of LRR domains in pattern recognition, humans might develop a pattern recognition protein for certain types of coronaviruses, in this case SARS-CoV-1 and -2. The robust and specific inhibitory effect of LRRC15 on multiple variants of SARS-CoV-2 and SARS-CoV-1 (but not MERS-CoV) suggests an arms race between humans and coronaviruses. While this manuscript was in preparation, LRRC15 was also identified as a SARS-CoV-2 spike binding factor in two preprints [43, 44]. This demonstrates the LRRC15-spike interaction is robust and detectable in multiple cell line models. Albeit the mechanism of viral restriction is different, several interferon-stimulated genes (ISGs) such as LY6E, CH25H, and IFITMs are known to restrict coronavirus entry by interfering with spike protein-mediated membrane fusion [45–47] or by interfering with endosome-mediated processes [48, 49]. Unlike these ISGs, LRRC15-mediated inhibition of viral entry is directly mediated by interaction with the spike, which resembles PAMP receptors. LRRC15 is not induced by interferons (**S6D Fig**). As LRRC15 redistributes adenovirus receptor (CAR) affecting the delivery of adenovirus to cells, it seems that LRRC15 may act as an antiviral factor via different modes [50]. It remains to be elucidated whether LRRC15 requires intracellular signaling through its cytosolic domain or requires other cellular proteins for the inhibition.

The *trans*-inhibition of viral entry by LRRC15 is striking. The co-culture of ACE2+LRRC15-cells and ACE2-LRRC15+ cells exhibited a significant suppression of viral infection with spike proteins of two different SARS-CoV-2 variants, indicating the antiviral effect of LRRC15 can be broad in a physiological context. During SARS-CoV-2 infection in the lung of human patients, LRRC15 may promote viral control by functioning as an entry inhibitor, likely in a non-cell-autonomous manner given that *ACE2* and *LRRC15* are expressed in distinct (but potentially neighboring) cell types. Thus, while the emergence of a *LRRC15*^+^ pathological fibroblast population may ultimately drive the fibrotic changes observed in patients with COVID-19, these fibroblasts may initially play a protective role by contributing to viral clearance during the acute phase of infection through their expression of LRRC15 on cell surface, subsequently paving the way for the transition towards tissue repair and remodeling.

In conclusion, this work reveals that LRRC15 is a receptor for SARS-CoV-2 spike and represents a pattern-recognition-like inhibition of viral entry by directly interacting with the spike protein. This study provides an insight into therapeutic development and a better understanding of COVID-19.

## MATERIALS AND METHODS

### Cell culture

HEK293T (#CRL-3216), HeLa (#CCL-2), and A375 (#CRL-1619) cells were purchased from ATCC. Cells were cultured in Dulbecco’s Modified Eagle Medium (DMEM, Gibco, #11995-081) supplemented with 10% fetal bovine serum (FBS) and 2.5% HEPES (Gibco, #15630-080) and detached using 0.05% trypsin-EDTA with phenol red (Gibco, #25300-120). After transfection, viral production media (DMEM with 10% FBS, 2.5% HEPES, and 1% bovine serum albumin) was used for lentiviral production. For A375 cells, 5 µg/mL blasticidin (Gibco, #A1113903) and 1 µg/mL puromycin (Gibco, #A1113803) were added as appropriate. For HeLa cells, 5 µg/mL blasticidin, 0.7 µg/mL puromycin, 200 or 400 µg/mL hygromycin (Gibco, #0-687-010), and 100 µg/mL zeocin (Thermo, #R25001) were added as appropriate.

### Generation of genetically modified cell lines

Individual sgRNAs (sgLRRC15 #1: GACATGCAGGCACTGCACTG; sgLRRC15 #2: AGTGTCAGCCCGGGACATGC; sgACE2: GTTACATATCTGTCCTCTCC) targeting the candidate genes were cloned into linearized pXPR_502 (Addgene, #96923) for CRISPR activation. Media was replaced with viral production media 12 hours after transfection. The supernatant was collected, spun at 4347 x g, and filtered using a 0.45 µm filter (Millipore) 36 hours after transfection. A375-dCas or HeLa-dCas cells were generated by transducing with pLenti-dCas9-VP64-Blast (Addgene, #61425). After 7 days of blasticidin selection, cells were transduced with 1 mL of harvested lentiviral stock and spun at 1200 x g for 90 minutes at 35°C. Cells were given 2 mL of fresh media and were incubated for 6 or 18 hours after spin transduction. Puromycin was used to select for successfully transduced cells.

Stable ACE2 expressing HeLa cells (HeLa-ACE2) were generated by transducing HeLa-dCas cells with pLENTI_hACE2_HygR (Addgene, #155296) followed selection with hygromycin. To generate a clonal cell line, transduced cells were plated in 96-well plates at single cell dilutions. After propagating the cells, each clone was screened for surface ACE2 expression by flow cytometry, and a clone with the highest ACE2 expression was selected and used for subsequent experiments.

For ectopic expression of LRRC15, a lentiviral vector pCDH-MSCV-T2A-Puro (System Biosciences, #CD522A-1) was modified to enable zeocin selection instead of puromycin. A codon-optimized LRRC15 ORF was cloned into pCDH-MSCV-T2A-Zeo with a C-terminal 3xFLAG tag and used to transduce HeLa-ACE2 followed by zeocin selection.

### CRISPR activation screen for SARS-CoV-2 spike binding

A list of 6011 surface proteins was obtained by integrating four datasets for plasma membrane proteins [51–54]. Four sgRNA sequences targeting each gene were picked from Calabrese genome-wide CRISPRa library [55] and 1,000 non-targeting control sgRNAs were included. The sgRNAs were cloned into pXPR_502 (Addgene, #96923) with assistance from the Genome Engineering and iPSC Center (GEiC) at Washington University in Saint Louis. 7.8 x 10^7^ A375-dCas cells were transduced with the CRISPRa library at ~0.3 MOI to make 2.4 x 10^7^ transduced cells, which is sufficient for the integration of each sgRNA into ~500 cells. At two days post transduction, puromycin was added and cells were selected for over a week.

For SARS-CoV-2 spike S1-Fc binding screen, 5 x 10^7^ cells per sample were washed with FACS buffer (1x PBS supplemented with 2% FBS and 1 mM EDTA) and incubated with 50 µg/mL SARS-CoV-2 spike S1 subunit-Fc fusion protein (R&D systems, #10623-CV-100) or human IgG1 isotype control (BioXCell, #BE0297) for 30 min at 4°C. After washing two times with FACS buffer, cells were stained with PE-conjugated anti-human IgG antibody (Southern Biotech, #9040-09) for 30 min at 4°C. Then the cells were washed two times with FACS buffer, fixed with 4% formaldehyde and subjected to sorting using FACSAriaIII (BD Biosciences). The top ~3% (fluorescence intensity) of the PE-positive cells were isolated. As a control, a same number of cells were stained with BV421 anti-hCD45 antibody (Biolegend, #368522) and the top 3% of the BV421-positive cells were sorted. Genomic DNA (gDNA) was extracted from the isolated cells and unsorted cells (“Input”) with QIAamp DNA Maxi kit (Qiagen, #51104).

### CRISPR screen sequencing and analysis

For Illumina sequencing, gDNA was used for PCR to amplify the integrated sgRNA sequences. PCR was performed in 96-well plates and each well containing up to 10 µg of gDNA in a total of 100 µL reaction mixture consisting of Titanium Taq DNA polymerase buffer and enzyme (Takara, #639209), deoxynucleoside triphosphate, dimethylsulfoxide (5%), P5 stagger primer mix (0.5 µM) and uniquely barcoded P7 primer (0.5 µM). Samples were amplified with following PCR cycles: an initial 5 min at 95°C; followed by 28 or 30 cycles of 95°C for 30 s, 59°C for 30 s, and 72°C for 20 s; followed by a final 10 min at 72°C. PCR products from two wells per sample were pooled and purified with AMPure XP beads according to the manufacturer’s protocol (Beckman Coulter, #MSPP-A63880). Samples were sequenced on a NextSeq550 sequencer (Illumina). After demultiplexing according to the barcode sequences, reads were mapped to a reference file of sgRNAs in the surface CRISPRa library using LibraryAligner (https://gitlab.com/buchserlab/library-aligner). We calculated the log-fold change of sgRNAs in each sample relative to the unsorted cells and calculated the hypergeometric distribution to determine p-values.

### Flow cytometry

For SARS-CoV-2 spike subunit binding assay, cells were washed once with HBSS containing 2% FBS and incubated with 50 µg/mL S1-Fc, 200 µg/mL RBD-Fc (Sino biological, #40592-V08H) or NTD-Fc (Sino biological, #40591-V49H) for 30 min at 4°C, followed by washing two times with HBSS with 2% FBS. Cells were incubated with PE anti-human IgG for another 30 min at 4°C, washed two times, and fixed with 4% formaldehyde for 15 min. Cells were washed once, resuspended in HBSS with 2% FBS and analyzed by flow cytometry using FACSCelesta (BD Biosciences). To measure the surface expression of ACE2 and LRRC15, cells were washed with FACS buffer (1x PBS supplemented with 2% FBS and 1 mM EDTA) and stained with goat anti-ACE2 (R&D Systems, #AF933) at a 1:50 dilution or rabbit anti-LRRC15 (abcam, #ab150376) at a 1:100 dilution for 30 min at 4°C. Then the cells were washed two times and resuspended in FACS buffer containing the secondary antibodies at a 1:1000 dilution: AF647-labeled donkey anti-goat IgG (Invitrogen, #A32849) or AF488-labeled goat anti-rabbit IgG (Invitrogen, #A32731). After 30 min incubation at 4°C, the cells were washed two times, fixed with 4% formaldehyde for 15 min and washed and resuspended in FACS buffer before analyzing by flow cytometry using FACSCelesta (BD Biosciences) or Cytek Aurora spectral analyzer (Cytek Biosciences). Data was analyzed with Flowjo software.

### ELISA binding assay

To investigate the binding of LRRC15 to SARS-CoV2 spike proteins, ELISA assays were performed on immobilized spike protein-Fc. To this aim, 96-well Immulon 2HB flat bottom plates (Thermo) were coated with 2 µg/mL spike proteins with C-terminal Histidine (BEI resources, NR-55438, NR-55311, NR-55310, NR-52724, NR-53769, and NR-55307, NR-55614, and NR-53589) at 4°C overnight, followed by 1-hr blocking buffer containing 1x HBSS (Gibco, #14025-092) and 2% FBS (VWR, #104B16). The plates were then incubated with either ACE2-Fc (Sino biological, #10108-H02H, starting concentration, 10 µg/mL), LRRC15-Fc (Sino biological, #15786-H02H, 100 µg/mL) or human IgG1 isotype (BioXCell, #BE0297) (as negative control) serially diluted 4-fold for 2 hours. The plates were then incubated with goat anti-human IgG Fc secondary antibody, HRP (Thermo, #A18817) at a 1:3000 dilution for 1 hour at room temperature. Next, TMB substrate (Thermo Scientific, #ENN301) was added to the plates and then quenched with stop solution (Thermo Scientific, #PIN600). Absorbance at 450 nm were recorded with a BioTek synergy HT microplate reader. Three washes were performed between every incubation using 1x HBSS with 0.05% Tween-20. GraphPad Prism 9 software was used to perform nonlinear regression curve-fitting analyses of binding data to estimate dissociation constants (K_D_).

To determine which SARS-CoV-2 spike protein region contributes to the binding of LRRC15, the ELISA was performed by coating 96 well plates with 2 µg/mL LRRC15-His (AcroBio, #LR5-H52H3) in coating buffer (BD, #51-2713KC) overnight at 4°C. Following blocking with 2% FBS, the 4-fold serially diluted SARS-CoV-2 spike RBD-Fc recombinant protein (Sino biological, #40592-V02H) and SARS-CoV-2 spike S1 NTD-Fc (Sino biological, #40591-V41H) were added and incubated for 2 hours at room temperature. After washing, the HRP labeled goat anti-human IgG Fc secondary antibody was added and incubated for 1 hour. Subsequently, TMB substrate was added, and the enzymatic reaction was stopped by adding stop solution. The signal was read at 450 nm.

To compare the binding affinity of LRRC15 and ACE2 with other coronaviruses, the 96-well plates were coated overnight at 4°C with 2 µg/mL LRRC15-His or ACE2-His (Sino biological, #10108-H08H). This was followed by blocking with 2% FBS for 1 h at room temperature. Then either rabbit Fc tagged-MERS-CoV spike/RBD protein fragment (Sino biological, #40071-V31B1) or SARS-CoV spike/RBD protein fragment (Sino biological, #40150-V31B2) were 4-fold serially diluted (starting concentration, 16 µg/mL) and added on the plate for 2-hour incubation. Later, goat anti-rabbit IgG-HRP conjugate (Southern Biotech, #4030-05) diluted (1:3000) in HBSS was used to detect the bound MERS or SARS-CoV RBD fragment. The reaction of HRP with TMB developed a colorimetric signal. The absorbance value was read at 450 nm after adding stop solution.

To determine whether LRRC15 binds to ACE2, 96-well plates were coated with either 2 µg/mL LRRC15-His, spike glycoprotein from SARS-CoV-2 with C-terminal histidine (BEI resources, NR-53589) or spike glycoprotein from MERS-CoV, England 1 with C-terminal histidine (BEI resources, NR-53591) and incubated overnight at 4°C. After blocking at room temperature for 1 h with 2% FBS in HBSS, the ELISA plates were washed, and 4-fold serially diluted ACE2-Fc (starting from 160 µg/mL) were added for a 2-hr incubation. HRP-conjugated goat anti-human IgG Fc secondary antibody was used to detect the bound ACE2.

### ELISA competition assay

The ACE2-Fc were serially diluted 4-fold starting from 40 µg/mL. Each dilution was mixed with different concentrations of LRRC15-His (0, 1250, 5000, and 20000 ng/mL). These sample series were transferred to the plate which coated with spike glycoprotein from SARS-CoV-2, Wuhan-Hu-1 with C-terminal histidine (BEI resources, NR-53947) at 4°C, overnight. After 2 hours incubation, the wells were treated with goat anti-human IgG Fc secondary antibody, HRP at a 1:3000 dilution for 1 hour at room temperature. Chromogenic development was generated with TMB substrate and quenched with stop solution. Optical density was measured in a BioTek synergy HT microplate reader.

### Pseudovirus production

VSV-dG pseudoviral particles were produced as previously described [31]. Briefly, 8 x 10^6^ HEK293T cells were plated in 10-cm tissue culture dishes and transfected using Lipofectamine2000 (Invitrogen) with plasmids encoding different CoV spike proteins or VSV-G protein. Expression vectors for SARS-CoV-2 Wuhan-Hu-1 (Addgene, #149539), SARS-CoV-2 B.1.167.2 (Addgene, #172320), SARS-CoV-2 B.1.1.7 (Addgene, #170451), SARS-CoV-2 B.1.351 (Addgene, #170449), SARS-CoV-2 P.1 (Addgene, #170450), SARS-CoV-1 (Addgene, #170447), MERS-CoV (Addgene, #170448) and VSV-G (Addgene, #12259) were used. At 24 h post transfection, cells were incubated with replication restricted rVSVΔG∗G-GFP virus (Kerafast, #EH1019-PM) at ~5 MOI for 1 h at 37°C, 5% CO_2_ and the media was replaced with complete media. Anti-VSV-G (Sigma, #MABF2337) was added at final concentration of 1 μg/mL to neutralize residual rVSVΔG∗G. At ~24 h post inoculation, viral supernatant was harvested, cell debris were removed by centrifuging for 10 min at 1320 x g, and stored at −80°C in small aliquots.

### Pseudovirus entry assay

Cells were plated at 1 x 10^5^ cells per well in 24-well plates or 2 x 10^4^ cells per well in 96-well plates. The following day, media was removed from the cells and 150 μL or 50 μL of pseudotyped VSV were added. After incubating 1 h at 37°C, 5% CO_2_, virus-containing media was removed and the cells were incubated in complete media for 20~24 h at 37°C, 5% CO_2_. Cells were washed once and resuspended with FACS buffer and GFP-positive cells were measured by flow cytometry using FACSCelesta (BD Biosciences). For neutralization assay, SARS-CoV-2 spike-pseudotyped VSV was preincubated with serial 3-fold dilutions of ACE2-Fc, LRRC15-Fc, or human IgG control for 1 h at 37°C, 5% CO_2_ before adding to the cells.

### Pseudovirus attachment assay

2 x 10^5^ cells were incubated with 150 μL of VSV pseudotyped with SARS-CoV-2 spike in microcentrifuge tubes for 1 h on ice. Then the cells were washed three times with chilled complete media to remove unattached viral particles. The cells were lysed in TRIzol and subjected to RNA extraction. The viral copies were measured by RT-qPCR with primers targeting VSV-N mRNA and normalized to the expression of Actin.

### Quantitative Reverse Transcription-PCR

As previously described [56], RNA extraction for cell samples was performed using TRI Reagent with a Direct-zol-96 RNA Kit (Zymo Research), following the manufacturer’s protocol. The ImProm-II reverse transcriptase system (Promega) was used with random hexamers and 5 µL of extracted RNA to synthesize cDNA. qPCR assays for VSV were performed using SYBR dye and primers targeting VSV-N (primer 1: 5’-TGTCTACCAAGGCCTCAAATC-3’; primer 2: 5’-GTGTTCTGCCCACTCTGTATAA-3’).

Predesigned PrimeTime qPCR assays (IDT) were used to quantify expression of human genes: *ACTB* (Hs.PT.39a.22214847), *LRRC15* (Hs.PT.58.26559170), *MX1* (Hs.PT.58.26787898), and *IFI44* (Hs.PT.58.21412074). A standard curve was used to determine absolute gene copy. qPCR results were normalized to the housekeeping gene *ACTB*.

### Analysis of human lung scRNA-seq datasets

Human lung scRNA-seq datasets from non-COVID-19 patients were accessed from the Human Lung Cell Atlas (https://github.com/krasnowlab/HLCA) (Synapse #syn21041850), and from the Tissue Stability Cell Atlas (https://www.tissuestabilitycellatlas.org/) (PRJEB31843). Preprocessed R objects were downloaded from the respective repositories and utilized for analysis of cell type-specific expression patterns using Seurat [57].

Human lung scRNA-seq datasets from deceased COVID-19 patients and non-COVID-19 controls were accessed from the Single Cell Portal of the Broad Institute. We downloaded the preprocessed data from Melms et al [36] (SCP1219) and Delorey et al [37] (SCP1052). For both datasets, we used Seurat to filter out all annotated doublets prior to investigating cell type-specific expression patterns. We also filtered out cells from non-COVID-19 patients for visualization of *ACE2* and *LRRC15* expression. For the Melms dataset, we also compared the relative proportions of each of the fibroblast subpopulations among total fibroblasts, using a two-tailed unpaired Mann-Whitney test to assess statistical significance.

### Analysis of bulk RNA-seq datasets

The Genotype-Tissue Expression (GTEx) project was supported by the Common Fund of the Office of the Director of the National Institutes of Health, and by NCI, NHGRI, NHLBI, NIDA, NIMH, and NINDS [35, 58]. Bulk RNA-seq normalized TPM matrices were accessed from the GTEx Portal (https://gtexportal.org/home/datasets) on March 18, 2020, release v8 and subsequently filtered to lung samples only. Gene expression data used in this study are publicly available on the web portal and have been de-identified. Detailed clinical annotations of the GTEx cohort were obtained as controlled access data through dbGaP (phs000424.v8.p2).

To test the association between various clinical factors and *LRRC15* expression in the GTEx cohort, we employed a similar approach as previously described [59], constructing a multivariable linear regression model using age, sex, diabetes (type or type 2), hypertension, body mass index (BMI), smoking, and ventilator status at time of death as predictor variables for log-transformed *LRRC15* expression. The resulting regression estimates were visualized as forest plots with 95% confidence intervals. To visualize *LRRC15* expression values after adjustment for all other clinical covariates except ventilator status, we summed the intercept and residuals from another multivariable linear regression model, omitting ventilator status as a predictor variable.

For analysis of genes correlated with *LRRC15* expression in the GTEx dataset, we computed the Spearman correlation between *LRRC15* and all other genes represented in the GTEx dataset. We performed multiple-hypothesis correction by the Benjamini-Hochberg method. For direct comparison of *ACE2* and *LRRC15* expression, we utilized log-transformed expression values to compute the linear regression line with 95% confidence intervals.

To compare *LRRC15* expression in the lungs of deceased COVID-19 patients with high vs low viral load [39], we accessed the raw count data from GSE150316 and performed differential expression analysis using DESeq2 [60]. We performed multiple-hypothesis correction by the Benjamini-Hochberg method.

To compare LRRC15 expression in lung cell lines infected with SARS-CoV-2 vs mock controls [61], we accessed the raw count data from GSE147507 and performed differential expression analysis using DESeq2 as above. For visualization purposes, we further extracted the normalized expression values for *LRRC15* in each of the samples and tested for statistical significance by two-tailed unpaired t-test.

### Trans-inhibition assay

4 x 10^3^ HeLa-ACE2 cells and 1.6 x 10^4^ HeLa or HeLa-sgLRRC15 cells (1:4 ratio) were co-plated per well in 96-well plates. For 1:1 ratio co-culture, 1 x 10^4^ HeLa-ACE2 cells and 1 x 10^4^ HeLa or HeLa-sgLRRC15 cells were plated per well. The following day, media was removed from the cells and 50 μL of pseudotyped VSV were added. After incubating 1 h at 37°C, 5% CO_2_, virus-containing media was removed and the cells were incubated in complete media for 20 h at 37°C, 5% CO_2_. Cells were washed once and resuspended with FACS buffer and GFP-positive cells were measured by flow cytometry using FACSCelesta (BD Biosciences). To compare GFP expressions in ACE2- and LRRC15-positive cells, pseudovirus-infected cells were stained for surface ACE2 and LRRC15 as described above with following secondary antibodies: AF405-labeled donkey anti-goat IgG (Invitrogen, #A48259) and PE-labeled donkey anti-rabbit IgG (Jackson ImmunoResearch, #711-116-152).

### Microscopic analysis

8 x 10^3^ HeLa-ACE2 cells and 3.2 x 10^4^ HeLa or HeLa-sgLRRC15 cells (1:4 ratio) were co-plated per well in 8-well chamber slides (Nunc). The following day, cells were inoculated with 75 μL of SARS-CoV-2 pseudovirus for 1 h on ice. Cells were washed three times with chilled media to remove unattached viral particles and placed back to 37°C, 5% CO_2_ to allow internalization. At 0, 30, or 60 min after internalization, cells were fixed by incubation of 4% paraformaldehyde in PBS for 15 min at room temperature and permeabilized with 0.1% Triton X-100 for 10 min. Cells were subsequently incubated with recombinant anti-LRRC15 antibody (Abcam, #ab150376) and SARS-CoV-2 spike antibody [1A9] (GeneTex, #GTX632604) at 1:100, followed by incubation with 1:500-diluted Alexa Fluor 555 conjugated goat anti-rabbit IgG antibody (Abcam, #ab150078) and 1:200-diluted Alexa Fluor 488 conjugated goat anti-mouse IgG antibody (Abcam, #ab150117) for 60 min at room temperature. The coverslips were mounted on a slide using prolong^TM^ glass antifade mountant with NucBlue^TM^ (Invitrogen, #P36981). The fluorescence images were recorded using a Zeiss microscope.

### Interferon stimulation

3 x 10^5^ A375 cells were plated per well in 24-well plates. The following day, Universal IFN-I (R&D systems, #11200-1) or IFN-λ2 (R&D systems, #8417-IL-025/CF), was added at 100 U/mL or 100 ng/mL, respectively, and incubated at 37°C, 5% CO_2_. After 6-h incubation, cells were harvested and subjected to RNA extraction and qPCR analysis as described above.

### Statistical analysis

Statistical significance was determined using GraphPad Prism 9 software. Experiments were analyzed by one-way or two-way ANOVA with Dunnett’s multiple comparisons test or unpaired two-pair t test as indicated.

## ACKNOWLEDGMENTS

We thank Robert Orchard for manuscript review and discussion. We also thank Megan Baldridge, Rachel Rodgers, Leran Wang, and William Buchser for helping to establish the bioinformatic analysis pipeline. We acknowledge the Brown University Flow Cytometry and Sorting Facility, the Genomics facility, the Leduc Bioimaging facility, and the Lentivirus Core for help with critical analysis. This study was supported by NIH grants R00 AI141683 (S.L.), K08 AI128043 (C.B.W), R01 AI148467 (C.B.W.), T32 GM007205 (R.D.C.), F30 CA250249 (R.D.C.) and P20 GM119943 (O.D.L.); the Smith Family Awards Program for Excellence in Biomedical Research (S.L.); a Burroughs Wellcome Fund Career Award for Medical Scientists (C.B.W.); the Ludwig Family Foundation (C.B.W.), the Mathers Charitable Foundation (C.B.W.); an Emergent Ventures fast grant (C.B.W).

## AUTHOR CONTRIBUTIONS

Conceptualization, J.S., S.L.; Methodology, J.S., R.D.C., L.Z., E-Y.S., S.L.; Validation, J.S., M.P-H., L.Z., S.A.L.; Investigation, J.S., S.L.; Resources, E-Y.S., O.D.L., C.B.W., S.L.; Writing – Original Draft, J.S., R.D.C., L.Z., S.A.L., S.L.; Writing – Review & Editing, M.P-H., E-Y.S., O.D.L., C.B.W.; Funding Acquisition, C.B.W., S.L.; Supervision, C.B.W., S.L.

