## Supplementary figures for "LRRC15 is an inhibitory receptor blocking SARS-CoV-2 spike-mediated entry *in trans*"

### S1 Fig.

**A**

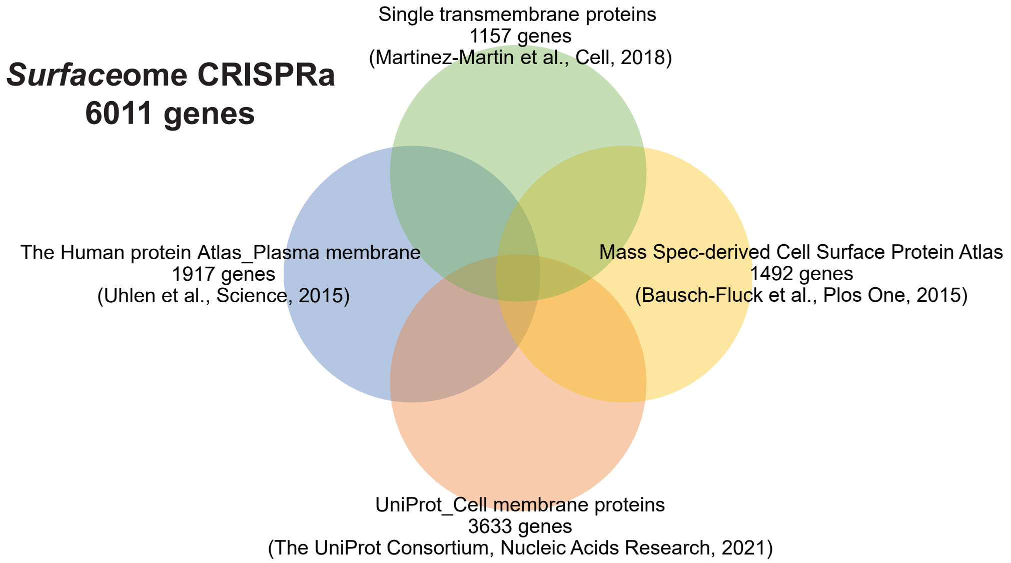


**B**

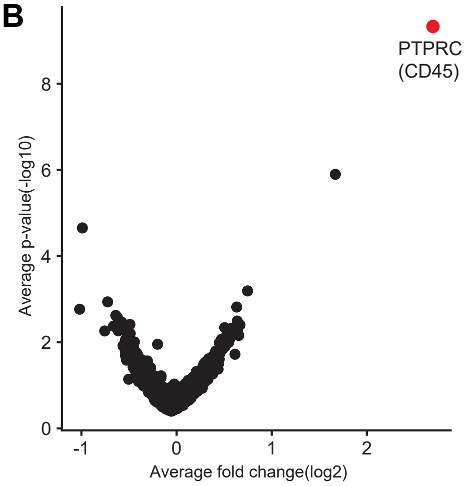


**S1 Fig. A focused CRISPR activation library was designed to induce surface proteins located on cellular plasma membrane**

(A) Venn diagram shows the composition of the surfaceome CRISPRa library.

(B) Volcano plot showing sgRNAs enriched or depleted in cells sorted after staining with anti-CD45.

### S2 Fig.

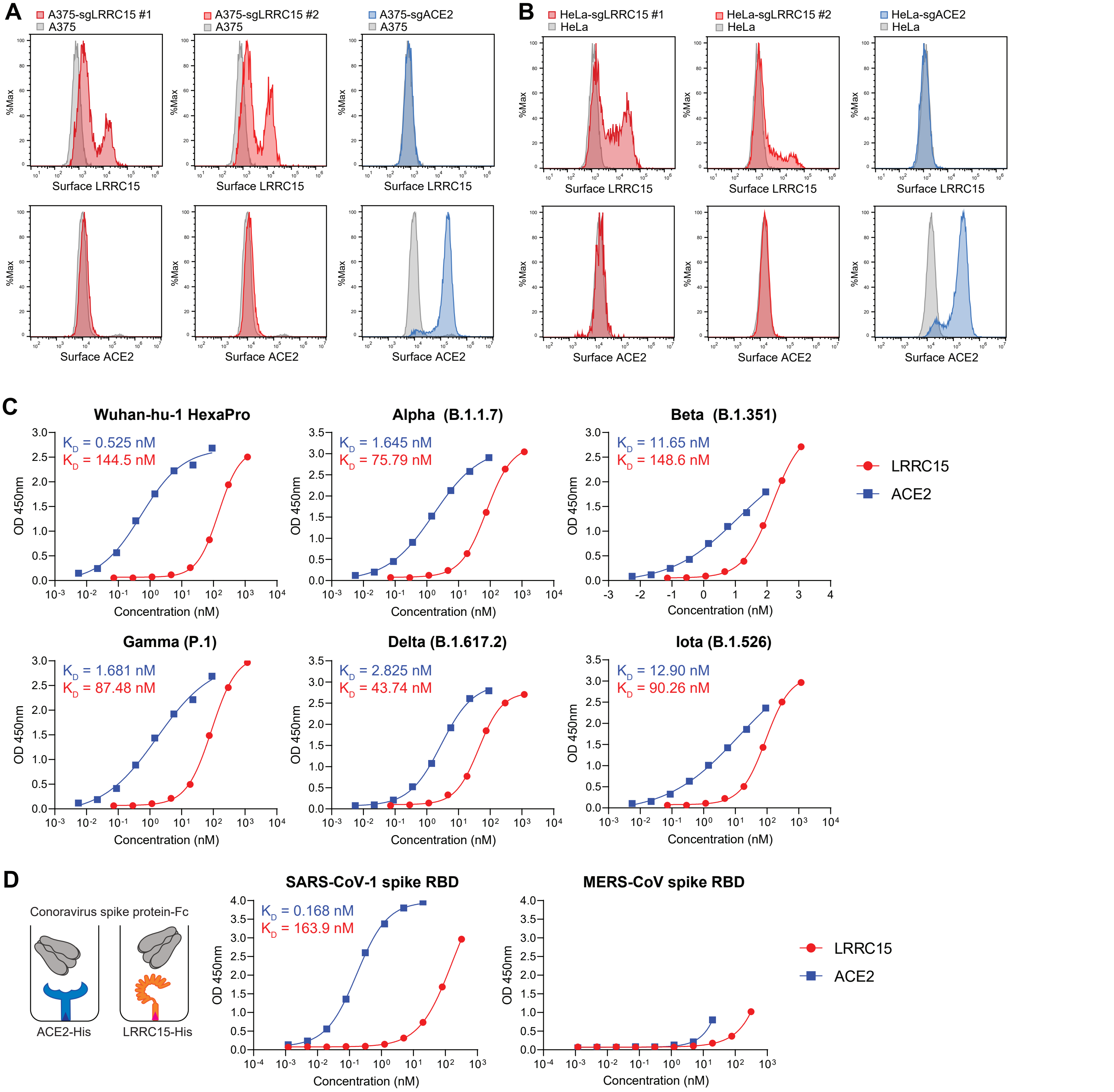


**S2 Fig. LRRC15 binds with spike proteins of SARS-CoV-2 variants and SARS-CoV-1**
(A-B) A375 cells (A) or HeLa cells (B) were transduced with indicated activating sgRNAs and surface expression of LRRC15 and ACE2 was measured by flow cytometry.
(C) The binding of ACE2 or LRRC15 to immobilized histidine-tagged spike proteins from SARS-CoV-2 mutant variants was measured using ELISA assay.
(D) The binding affinity of LRRC15 to MERS-CoV spike protein RBD fragment and SARS-CoV spike protein RBD fragment was determined by ELISA.

### S3 Fig.

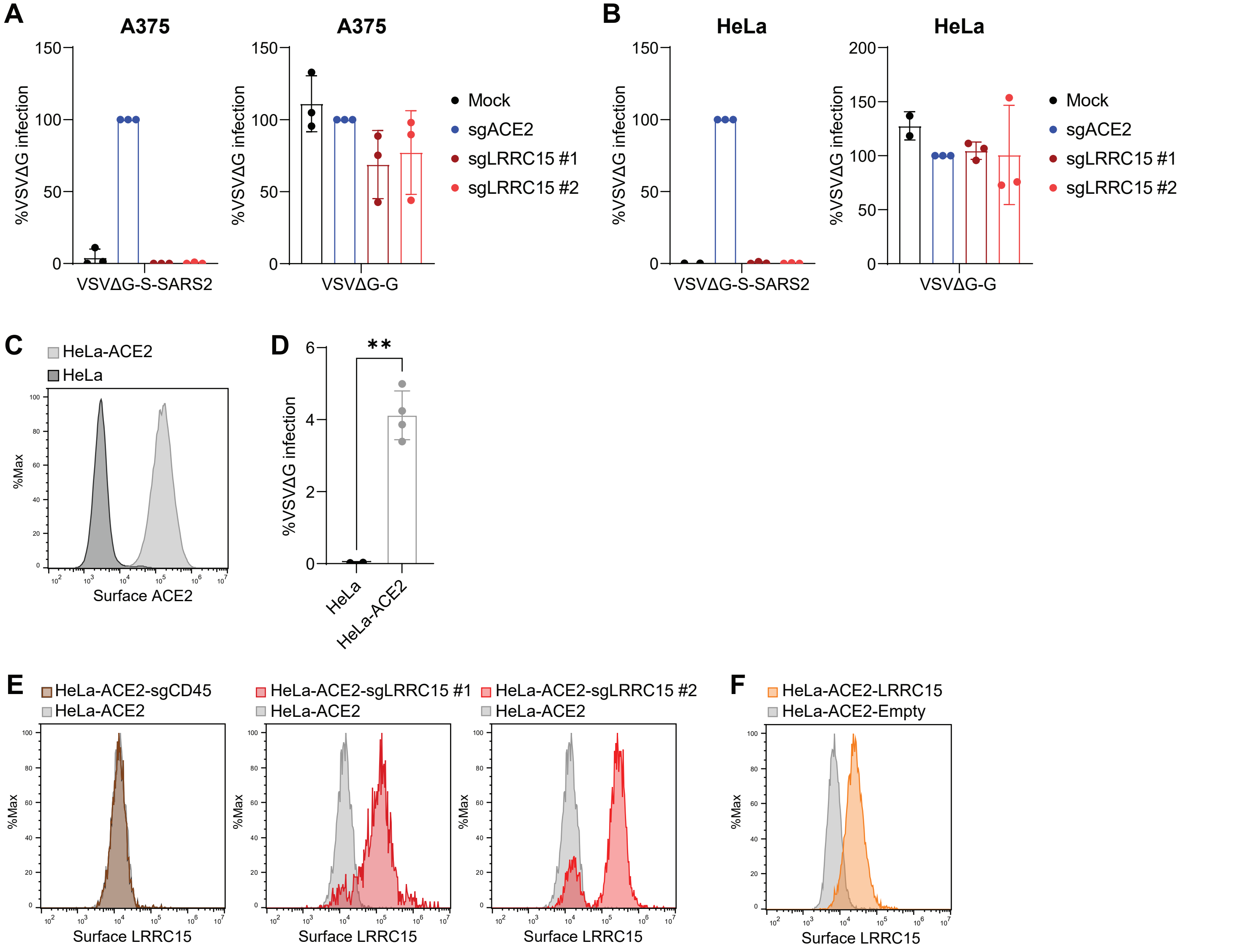


**S3 Fig. LRRC15 is not an entry receptor for SARS-CoV-2**
(A-B) A375 cells (A) or HeLa cells (B) transduced with indicated activating sgRNAs were infected with VSV pseudoviruses VSVΔG-S-SARS2 or VSVΔG-G. GFP signal was measured at 20 hpi by flow cytometry and normalized to sgACE2-transduced cells (n=3).
(C) HeLa cells stably expressing ACE2 (HeLa-ACE2) were assessed for cell surface expression of ACE2 by flow cytometry.
(D) HeLa or HeLa-ACE2 cells were infected with VSV pseudovirus harboring SARS-CoV-2 spike and GFP signal was measured at 20 hpi by flow cytometry (n=4).
(E) HeLa-ACE2 cells transduced with indicated activating sgRNAs were assessed for surface expression of LRRC15 was measured by flow cytometry.
(F) HeLa-ACE2 cells expressing LRRC15 or empty vector were assessed for surface expression of LRRC15 was measured by flow cytometry.
Data represent means ± SEM (A, B, D). Data were analyzed unpaired two-tailed t test (D); **p < 0.01.

### S4 Fig.

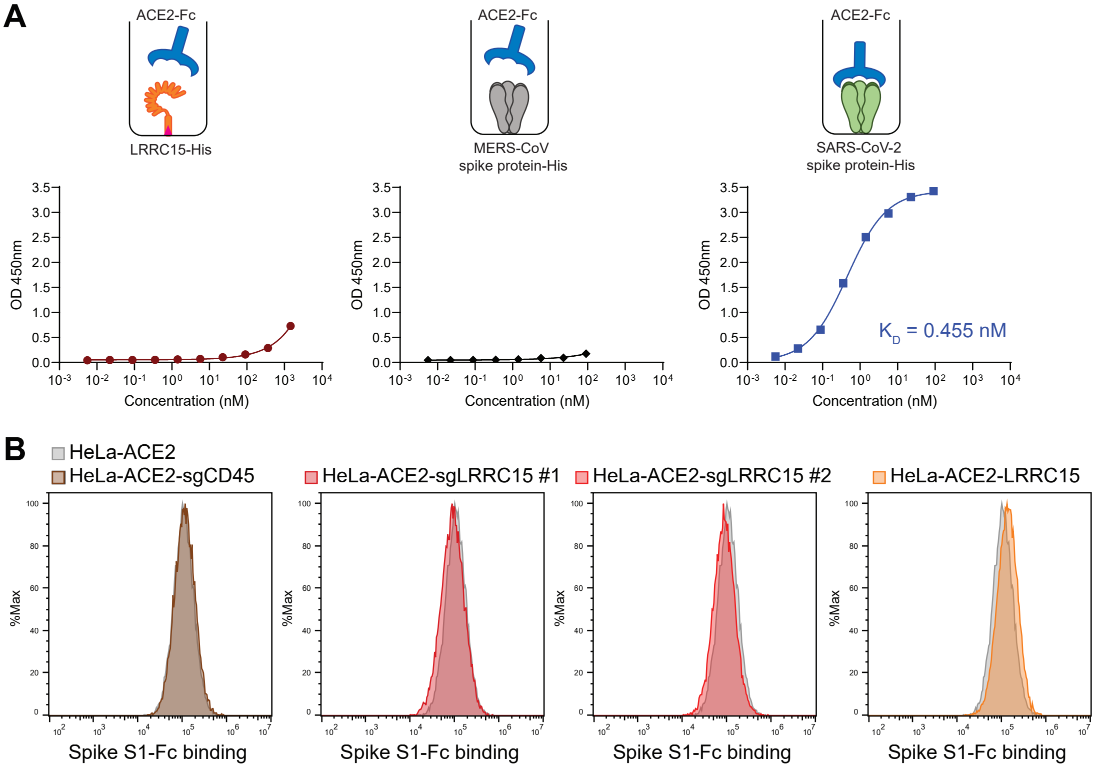


**S4 Fig. LRRC15 does not interact with ACE2**

(A) Binding of Fc-tagged recombinant human ACE2 to His-tagged LRRC15 was measured by ELISA. SARS-CoV-2 spike protein was used as a positive control, MERS-spike protein as a negative control.

(B) Binding of SARS-CoV-2 spike S1-Fc to HeLa-ACE2 cells transduced with indicated activating sgRNAs or a LRRC15-expressing vector was measured by flow cytometry.

### S5 Fig.

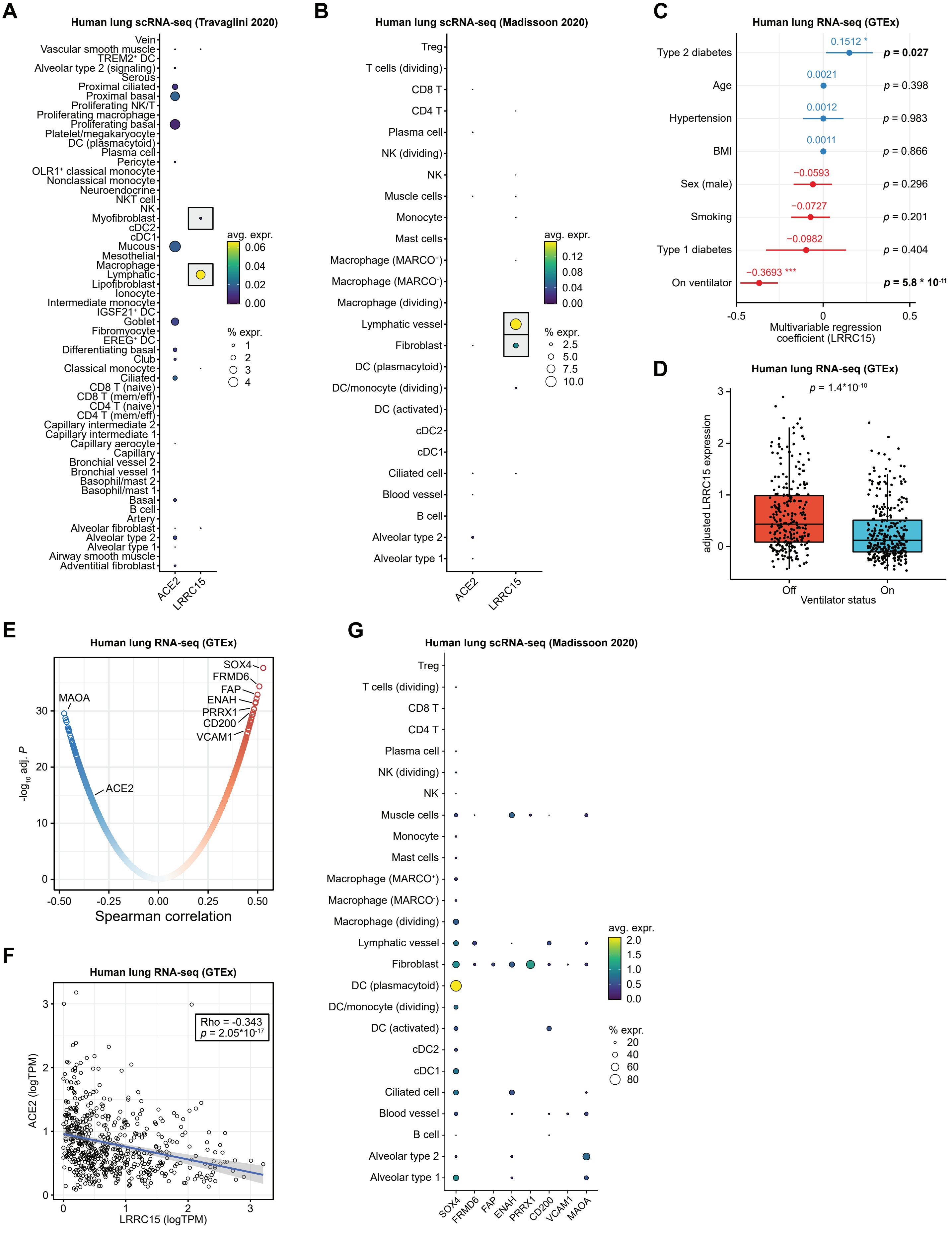
A
Human lung scRNA-seq (Travaglini 2020)
Vein
Vascular smooth muscle
TREM2⁺ DC
Alveolar type 2 (signaling)
Serous
Proximal ciliated
Proximal basal
Proliferating NK/T
Proliferating macrophage
Proliferating basal
Platelet/megakaryocyte
DC (plasmacytoid)
Plasma cell
Pericyte
OLR1⁺ classical monocyte
Nonclassical monocyte
Neuroendocrine
NKT cell
NK
Myofibroblast
cDC2
cDC1
Mucous
Mesothelial
Macrophage
Lymphatic
Lipofibroblast
Ionocyte
Intermediate monocyte
IGSF21⁺ DC
Goblet
Fibromyocyte
EREG⁺ DC
Differentiating basal
Club
Classical monocyte
Ciliated
CD8 T (naive)
CD8 T (mem/eff)
CD4 T (naive)
CD4 T (mem/eff)
Capillary intermediate 2
Capillary intermediate 1
Capillary aerocyte
Capillary
Bronchial vessel 2
Bronchial vessel 1
Basophil/mast 2
Basophil/mast 1
Basal
B cell
Artery
Alveolar fibroblast
Alveolar type 2
Alveolar type 1
Airway smooth muscle
Adventitial fibroblast
ACE2
LRRC15
avg. expr.
0.06
0.04
0.02
0.00
% expr.
1
2
3
4
B
Human lung scRNA-seq (Madissoon 2020)
Treg
T cells (dividing)
CD8 T
CD4 T
Plasma cell
NK (dividing)
NK
Muscle cells
Monocyte
Mast cells
Macrophage (MARCO⁺)
Macrophage (MARCO⁻)
Macrophage (dividing)
Lymphatic vessel
Fibroblast
DC (plasmacytoid)
DC/monocyte (dividing)
DC (activated)
cDC2
cDC1
Ciliated cell
Blood vessel
B cell
Alveolar type 2
Alveolar type 1
ACE2
LRRC15
avg. expr.
0.12
0.08
0.04
0.00
% expr.
2.5
5.0
7.5
10.0
C
Human lung RNA-seq (GTEx)
Type 2 diabetes
0.1512 *
p = 0.027
Age
0.0021
p = 0.398
Hypertension
0.0012
p = 0.983
BMI
0.0011
p = 0.866
Sex (male)
−0.0593
p = 0.296
Smoking
−0.0727
p = 0.201
Type 1 diabetes
−0.0982
p = 0.404
On ventilator
−0.3693 ***
p = 5.8 * 10⁻¹¹
-0.5
0
0.5
Multivariable regression
coefficient (LRRC15)
D
Human lung RNA-seq (GTEx)
p = 1.4*10⁻¹⁰
adjusted LRRC15 expression
3
2
1
0
Off
On
Ventilator status
E
Human lung RNA-seq (GTEx)
SOX4
FRMD6
FAP
ENAH
PRRX1
CD200
VCAM1
MAOA
ACE2
-log₁₀ adj. P
30
20
10
0
-0.50
-0.25
0.00
0.25
0.50
Spearman correlation
F
Human lung RNA-seq (GTEx)
Rho = -0.343
p = 2.05*10⁻¹⁷
ACE2 (logTPM)
3
2
1
0
LRRC15 (logTPM)
G
Human lung scRNA-seq (Madissoon 2020)
Treg
T cells (dividing)
CD8 T
CD4 T
Plasma cell
NK (dividing)
NK
Muscle cells
Monocyte
Mast cells
Macrophage (MARCO⁺)
Macrophage (MARCO⁻)
Macrophage (dividing)
Lymphatic vessel
Fibroblast
DC (plasmacytoid)
DC/monocyte (dividing)
DC (activated)
cDC2
cDC1
Ciliated cell
Blood vessel
B cell
Alveolar type 2
Alveolar type 1
SOX4
FRMD6
FAP
ENAH
PRRX1
CD200
VCAM1
MAOA
avg. expr.
2.0
1.5
1.0
0.5
0.0
% expr.
20
40
60
80

**S5 Fig. LRRC15 expression in the human lung is cell type-specific and inversely associated with ventilator usage.**
(A-B) Cell type-specific expression of ACE2 and LRRC15, assessed by scRNA-seq of human lungs from (A) the Human Lung Cell Atlas [33] or (B) the Tissue Stability Cell Atlas [34].
(C) Forest plot of the multivariable regression coefficients for various clinical features as predictors for lung expression of LRRC15 (from GTEx; n = 554 samples) [35].
(D) LRRC15 expression after adjustment for clinical features noted in (C), classifying each lung sample by ventilator status at the time of death. Statistical significance was assessed by two-tailed unpaired Mann-Whitney test.
(E) Spearman correlation analysis between LRRC15 and all other genes, based on RNA-seq of human lung samples from GTEx. Multiple hypothesis correction was performed by the Benjamini-Hochberg method.
(F) Scatter plot comparing log normalized expression of LRRC15 and ACE2 across human lung samples. Spearman rho = -0.343, p = 2.05*10-17. The linear regression line is overlaid, with 95% confidence intervals shaded in.
(G) Cell type-specific expression of genes that are highly correlated with LRRC15, assessed by scRNA-seq of human lungs from the Tissue Stability Cell Atlas.

**S6 Fig.**

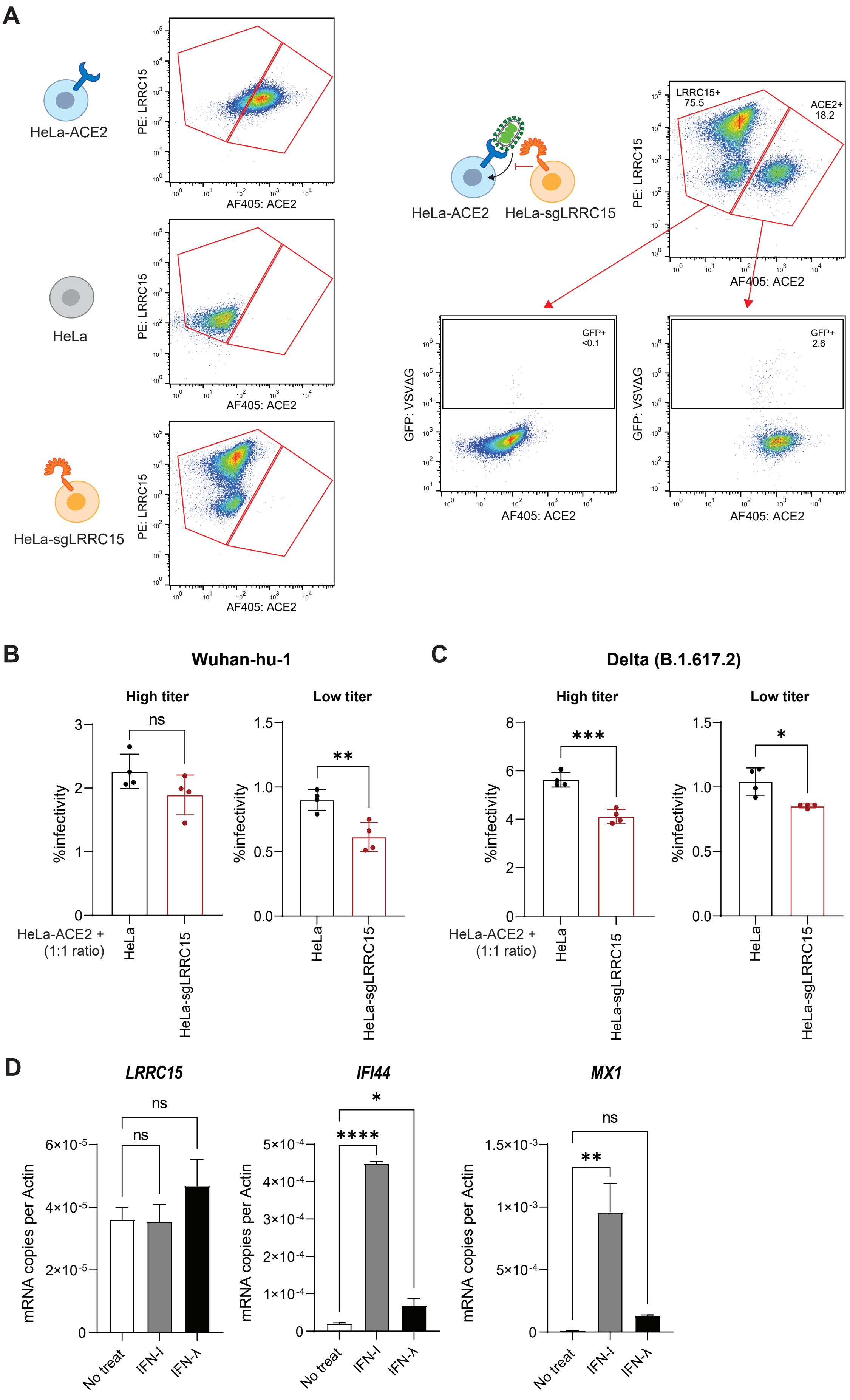
A
HeLa-ACE2
HeLa
HeLa-sgLRRC15
PE: LRRC15
AF405: ACE2
HeLa-ACE2 HeLa-sgLRRC15
LRRC15+
75.5
ACE2+
18.2
GFP+
<0.1
GFP+
2.6
GFP: VSVΔG
B
Wuhan-hu-1
C
Delta (B.1.617.2)
High titer
Low titer
%infectivity
ns
**
***
*
HeLa-ACE2 +
(1:1 ratio)
HeLa
HeLa-sgLRRC15
D
LRRC15
IFI44
MX1
mRNA copies per Actin
****
No treat
IFN-I
IFN-λ

**S6 Fig. LRRC15 inhibits ACE2-mediated SARS-CoV-2 entry in trans**

(A) Left, surface ACE2 and LRRC15 expressions of HeLa, HeLa-ACE2, or HeLa-sgLRRC15 cells were measured by flow cytometry. Right, after co-culturing HeLa-ACE2 cells with HeLa-sgLRRC15 cells and infecting with VSVΔG-S-SARS2, surface ACE2 and LRRC15 expressions were measured by flow cytometry. GFP expressions were compared in LRRC15+ (HeLa-sgLRRC15) cells and ACE2+ (HeLa-ACE2) cells.

(B) HeLa-ACE2 cells were co-cultured with HeLa or HeLa-sgLRRC15 cells at 1:1 ratio and infected with VSV pseudovirus harboring spike of SARS-CoV-2 Wuhan-hu-1 strain at high or low titer. GFP signal was measured at 20 hpi by flow cytometry (n=4). Representative of three independent experiments are shown.

(C) Trans-inhibition assay was performed as in (B) with VSV pseudovirus harboring spike of SARS-CoV-2 Delta variant (B.1.617.2 strain) (n=4). Representative of three independent experiments are shown.

(D) A375 cells were treated with Universal IFN-I (100 U/mL) or IFN-λ2 (100 ng/mL) and harvested after 6 h. RNA expressions of LRRC15, IFI44, and MX1 was measured by qPCR and normalized to ACTB.

Data represent means ± SEM and were analyzed by unpaired two-tailed t test (B, C) or one-way ANOVA with Dunnett's multiple comparisons test (D). ns, not significant; $*p < 0.05$; $**p < 0.01$; $***p < 0.001$; $****p < 0.0001$.
